# Interactions between Nodal and Wnt signalling Drive Robust Symmetry-Breaking and Axial Organisation in *Gastruloids* (Embryonic Organoids)

**DOI:** 10.1101/051722

**Authors:** D.A. Turner, C.R. Glodowski, L. Alonso-Crisostomo, P. Baillie-Johnson, P.C. Hayward, J. Collignon, C. Gustavsen, P. Serup, C. Schröter, A. Martinez Arias

## Abstract

Generation of asymmetry within the early embryo is a critical step in the establishment of the three body axes, providing a reference for the patterning of the organism. To study the establishment of asymmetry and the development of the anteroposterior axis (AP) in culture, we utilised our ‘*Gastruloid*’ model system. ‘*Gastruloids*’, highly reproducible embryonic organoids formed from aggregates of mouse embryonic stem cells, display symmetry-breaking, polarised gene expression and axial development, mirroring the processes on a time-scale similar to that of the mouse embyro. Using *Gastruloids* formed from mouse ESCs containing reporters for Wnt, FGF and Nodal signalling, we were able to quantitatively assess the contribution of these signalling pathways to the establishment of asymmetry through single time-point and live-cell fluorescence microscopy.

During the first 24-48h of culture, interactions between the Wnt/*β*-Catenin and Nodal/TGF/*β* signalling pathways promote the initial symmetry-breaking event, manifested through polarised *Brachyury* (T/Bra) expression. Neither BMP nor FGF signalling is required for the establishment of asymmetry, however Wnt signalling is essential for the amplification and stability of the initial patterning event. Additionally, low, endogenous levels of FGF (24-48h) has a role in the amplification of the established pattern at later time-points.

Our results confirm that *Gastruloids* behave like epiblast cells in the embryo, leading us to translate the processes and signalling involved in pattern formation of *Gastruloids* in culture to the development of the embryo, firmly establishing *Gastruloids* as a highly reproducible, robust model system for studying cell fate decisions and early pattern formation in culture.

## 1 Introduction

The emergence of the anteroposterior (AP) axis during the early stages of animal development is a fundamental patterning event that guides the spatial organisation of tissues and organs during embryogenesis. Comparative studies reveal that, although this process differs from one organism to another, in all cases it involves a symmetry-breaking event within a molecular or cellular isotropic system that results in the asymmetric localisation of signalling centres that will drive subsequent patterning events. Dipteran (e.g *Drosophila*) and amniote (e.g chicken and mouse) embryos provide extreme examples of the strategies associated with these processes. In *Drosophila*, the symmetry is broken within a single cell, the oocyte, which acquires both the AP and dorsoventral (DV) axes through well characterised processes of RNA and protein localisation that then serve as references for the rapid patterning of the embryo after fertilisation [1, 2]. On the other hand, in birds and mammals the process occurs in the developing embryo, within a multicellular system, and leads to the assignation of fates to cell populations [3–7].

Efforts to understand the mechanisms that pattern early embryos have commonly relied on genetic approaches: identification of mutations that disrupt the process of the genes affected in the mutations and, guided by the molecular nature of the protein encoded by those genes, the assignation of molecular events to the process [9, 10]; this approach has been successful. However, while it allows some insights into the processes at work, it has limitations. It reveals what is necessary but not what is sufficient, it can conflate correlation and causation, has problems with redundancies and cannot easily probe for the role of mechanical forces, which experiments suggest also play a role in the early events [11, 12]. A complementary approach requires an experimental system that allows a precise and reliable perturbation of the patterning process, the quantitative analysis of the experimental results and, if possible, the exploitation of available genetics. Importantly, such system should mimic the embryo.

We have established a non adherent culture system for mouse Embryonic Stem Cells (ESCs) in which small aggregates of cells undergo symmetry-breaking, polarisation of gene expression and axial development in a reproducible manner that mirrors events in embryos [13–15]. These polarised aggregates, that we call *Gastruloids*, provide a versatile and useful system that, in combination with genetics, fulfils many the requirements for experimental manipulations to study pattern formation in cell ensembles.

Here we use the *Gastruloid* system to study the signalling events that lead to their early patterning: the symmetry-breaking event and the polarised expression of Brachyury (*T/Bra*) that, in the embryo, reflects the establishment of the AP axis and presages the process of gastrulation. We find that during 24-48h of culture, interactions between Nodal and Wnt signalling promote an intrinsic symmetry-breaking event that is recorded in the expression of *T/Bra*; BMP signalling is not required for these events. These results confirm that *Gastruloids* behave like cells in the epiblast [13] and lead us to suggest that a similar spontaneous symmetry-breaking event occurs in the embryo where biases from the extraembryonic tissues, ensure its reproducible location next to the Extraembryonic Ectoderm. We also show that Wnt signalling plays an important role in the amplification and stabilisation of these events.

Our results establish *Gastruloids* as a robust experimental system to analyse mechanisms of fate decisions and pattern formation in mammals and provide some insights about the interactions between signalling pathways in the patterning of embryos, in particular about the role of epithelia and the extraembryonic tissues.

## 2 Materials and Methods

### Cell lines and routine cell culture

Bra::GFP [16], Nodal^*condH BE::Y F P*^ [17], AR8::mCherry (Smad2/3 reporter) [18], IBRE4-TA-Cerulean (BMP reporter) [18], Spry4::H2B-Venus (see below) and TCF/LEF::mCherry (TLC2) [19, 20] were cultured in serum supplemented with LIF and fœtal bovine serum (ESL medium) on gelatinised tissue-culture flasks and passaged every second day as previously described [14, 19, 21–23]. If cells were not being passaged, half the medium in the tissue culture flask was replaced with ESL. All cell lines were routinely tested and confirmed free from mycoplasma.

### Generation of Cell Lines

The Spry4::H2B-Venus reporter cell line was generated by combining *knockout first* targeting arms of the EUCOMM project [24] with a H2B-Venus reporter cassette and a neomycin resistance gene driven from a human *β*-actin promoter. This construct was integrated by homologous recombination into a cell line carrying a doxycycline-inducible Gata4-mCherry construct described in [25]. This cell contains a constitutively active Cerulean fluorescent protein to aid in image analysis [25]. See supplemental materials and methods for a detailed description of the generation of this cell line.

### Immunofluorescence and Microscopy

*Gastruloids* from the Nodal^condH B E::Y F P^ [17] cell line were fixed and stained for YFP, Brachyury and Nanog according to the protocol previously described [15]. Hoechst3342 was used to mark the nuclei (see Table S1 for the antibodies used and their dilutions). Confocal z-stacks of *Gastruloids* were generated using an LSM700 (Zeiss) on a Zeiss Axiovert 200 M using a 40 EC Plan-NeoFluar 1.3 NA DIC oil-immersion objective. Hoechst3342, Alexa-488, 568 and 633 were sequentially excited with 405, 488, 555 and 639 nm diode lasers respectively as previously described [23]. Data capture was carried out using Zen2010 v6 (Carl Zeiss Microscopy Ltd, Cambridge UK) and analysis of the average pixel intensities within each nucleus within a given z-plane was performed in Fiji [26]. The z-stacks were acquired of at least 4 *Gastruloids* per condition with a z-interval of 2.11μm for a maximum of 42.2μm.

Widefield, single-time point images of *Gastruloids* were acquired using a Zeiss AxioObserver.Z1 (Carl Zeiss, UK) in a humidified CO_2_ incubator (5% CO_2_, 37^°^C) with a 20x LD Plan-Neofluar 0.4 NA Ph2 objective with the correction collar set to image through plastic. Illumination was provided by an LED white-light system (Laser2000, Kettering, UK) in combination with filter cubes GFP-1828A-ZHE (Semrock, NY, USA), YFP-2427B-ZHE (Semrock, NY, USA) and Filter Set 45 (Carl Zeiss Microscopy Ltd. Cambridge, UK) used for GFP, YFP and RFP respectively and emitted light recorded using a back-illuminated iXon888 Ultra EMCCD (Andor, UK). Images were analysed using the ImageJ image processing package Fiji [26] and plugins therein as previously described [15]. Briefly, the florescence intensity was measured by a line of interest (LOI) drawn from the posterior to anterior region of the *Gastruloid* with the LOI width set to half the diameter of the *Gastruloid* at 48h (100px with the 20x objective). The background for each position was measured and subtracted from the fluorescence for each Gastruloid. Shape-descriptors were generated by converting brightfield images of *Gastruloids* to binary images and measuring them by particle detection.

Fluorescence levels were normalised to the maximum obtained in following Chi stimulation, and the maximum length of each *Gastruloid* was rescaled 1 unit. Average fluorescence traces of *Gastruloids* ± S.D. are shown.

For live imaging experiments, each well of a 96-well plate containing individual *Gastruloids* was imaged as described above using both the 20× (24-72h) and the 10× (72-96h) objectives described above, and images captured every 30 min for a maximum of 96h. All images were analysed in Fiji [26] using the LOI interpolator [27] with the LOI set to a width of 100px.

### Gastruloid culture and application of specific signals

Aggregates of mouse ESCs were generated as previously described [13, 15]. Briefly, mouse ESCs suspended in 40μl droplets of N2B27 were plated in round-bottomed low-adhesion 96-well plates and left undisturbed for 48h, following which, they were competent to respond to specific signals. The number of cells within each droplet was optimised to ensure that the size of the *Gastruloid* was ~150μm in diameter at 48h; this generally consisted of 400 cells/40μl/well (See Table S2 for the numbers of cells required for each cell line used in this study). In experiments which required the addition of specific factors to *Gastruloids* on the second day of aggregation (24-48h), 20μl medium was carefully removed with a multichannel pipette, and 20μl of N2B27 containing twice the concentration of factors was added. This method was preferable to the addition of smaller volumes containing higher concentrations of agonist/antagonists, as the data from these experiments showed more variation between *Gastruloids* (DAT, PB-J, AMA unpublished). Control experiments (e.g. DMSO or N2B27 only) showed that replacement of half the medium at this stage did not significantly alter the ability of *Gastruloids* to respond to signals on the third day. The next day, 150μl fresh N2B27 was added to each of the wells with a multichannel pipette for 1h to wash the *Gastruloids*; a time delay ensured that sample loss was prevented. Following washing, 150μl N2B27 containing the required factors was then applied. The small molecules used in this study and their concentrations are described in Table S3.

### Quantitative RT-PCR

*Gastruloids* (*n* = 64 per time-point) from T/Bra::GFP mouse ESCs, subjected to a Chi or DMSO pulse (between 48 and 72h AA), harvested at 48 or 72h AA, trypsinised, pelleted and RNA extracted using the RNeasy Mini kit (Qiagen, 74104) according to the manufacturers instruction as previously described [22]. Samples were normalised to the housekeeping gene *PPIA*. The primers for *BMP4, Cerl, Chordin, DKK, FGF4, FGF5, FGF8, Lefty1, Nodal, Ppia, Spry4, Wnt3* and *Wnt3a* are described in Table S4.

### Orientation of Gastruloids

To define the AP orientation of *Gastruloids*, we have assigned the point of T/Bra::GFP expression as the ‘Posterior’, as the primitive streak, which forms in the posterior of embryo, is the site of *T/Bra* expression in the embryo [28–30].

## 3 Results

### 3.1 Intrinsic Patterning in the Gastru-loids

As a reference for the evaluation and interpretation of our results, a summary of the main events that lead to the establishment of the anteroposterior (AP) axis in the mouse embryo is provided in **Fig. 1**. This process confines the start of gastrulation to a cell population in the proximal posterior edge of the embryo, next to the Extraembryonic Ectoderm, through a sequence of carefully choreographed interactions between extraembry-onic and embryonic tissues mediated by Nodal, BMP and Wnt signalling. The relationship between these three signals has been established in genetic experiments and places Nodal and Wnt signalling in the epiblast as the effective target of the process [31, 32].

**Figure 1:**
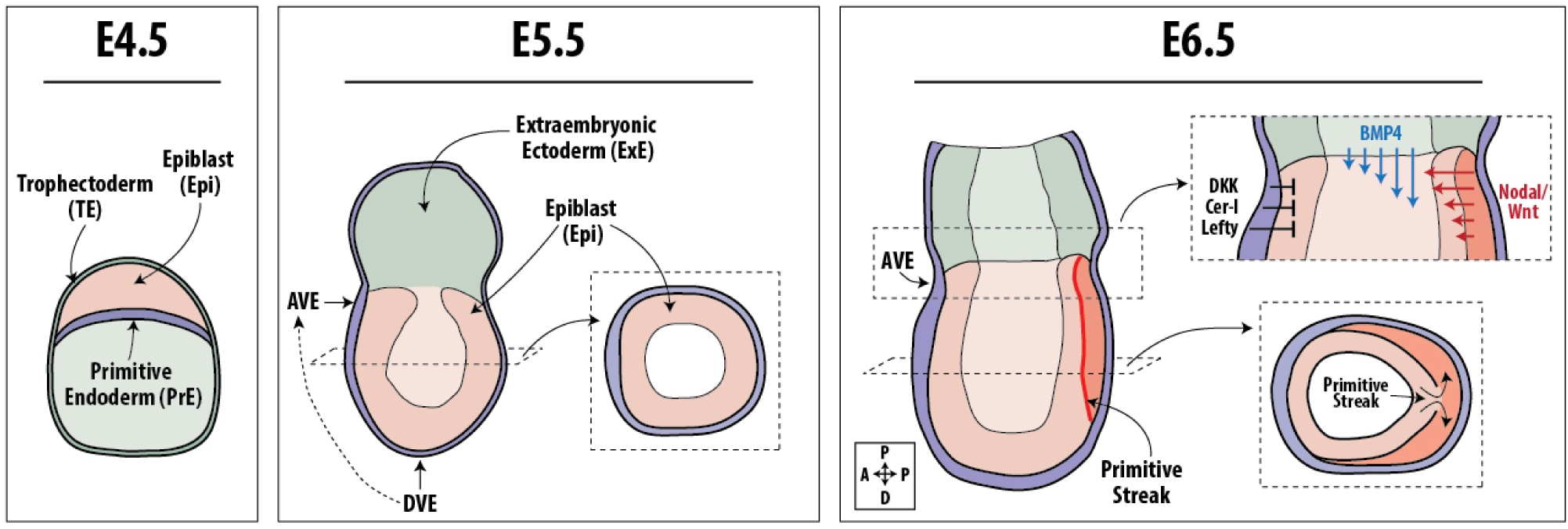
Establishment of the Anteroposterior Axis in the Mouse. At E4.5, following several rounds of cell division and the segregation of the extraembryonic tissues (the Primitive Endoderm (PrE) and the Trophoec-toderm (TE)), the embryo is represented in the blastocyst, a mass of equivalent cells that undergoes a process of epithelialisation. By stage E5.5 the embryo forms a radially symmetric cup-shaped epithelium, the Epiblast (Epi) containing around 300 cells that is attached to the Extra embryonic Ectoderm (ExE) on the proximal side and embedded in another epithelium, the Visceral Endoderm (VE) a derivative of the PrE. At this time, a symmetry-breaking event defines a group of cells within the VE that move to one side and give rise to the Anterior Visceral Endoderm (AVE), a cellular collective that defines the anterior region of the embryo on the adjacent Epi cells and, as a result, establishes the anteroposterior (AP) axis of the embryo. At this stage, the Epi experiences widespread BMP, Nodal and Wnt signalling but the AVE, acting as a source of antagonists of these signalling pathways (*Cerl, Lefty1* and *Dkk1*), restricts their effects to the side of the epithelium opposite to the AVE where they induce the localised expression of T/Bra, the start of gastrulation by initiation of the Primitive Streak and axial extension at the posterior end. Figure part adapted from [8].

To investigate initial patterning mechanisms during development, and to circumvent the difficulties involved in experimentation with embryos at this stage of development, we utilised our non-adherent culture method [13–15]. We have reported before that in this system, small numbers of ESCs upon specific culture treatment will undergo symmetry-breaking and axial elongation; here we confirm this. Using a T/Bra::GFP line that identifies the Bra-expressing cells as the posterior of the *Gastruloids*, we set out to study the mechanisms involved in the patterning of *Gastruloids*. (See Materials and Methods). After the initial aggregation period, *Gastruloid* cultures were kept individually for 2 days in N2B27 (**Fig. 2A**). If they were maintained in N2B27 for a further 72h (120h AA), a variety of patterns were observed with a large degree of inter-experimental variation. In general about 50% of individual *Gastruloids* exhibited different degrees of axial elongation and T/Bra::GFP polarisation while the rest were very varied. Following stimulation by signals such as the Wnt/*β*-Catenin agonist (Chi) between 48-72h AA, over 85% of the *Gastruloids* underwent axial elongation. Our results suggest that there is an intrinsic symmetry-breaking event between 24 and 48h AA and that exposure to Wnt/*β*-Catenin signalling plays a role in its stabilisation.

**Figure 2:**
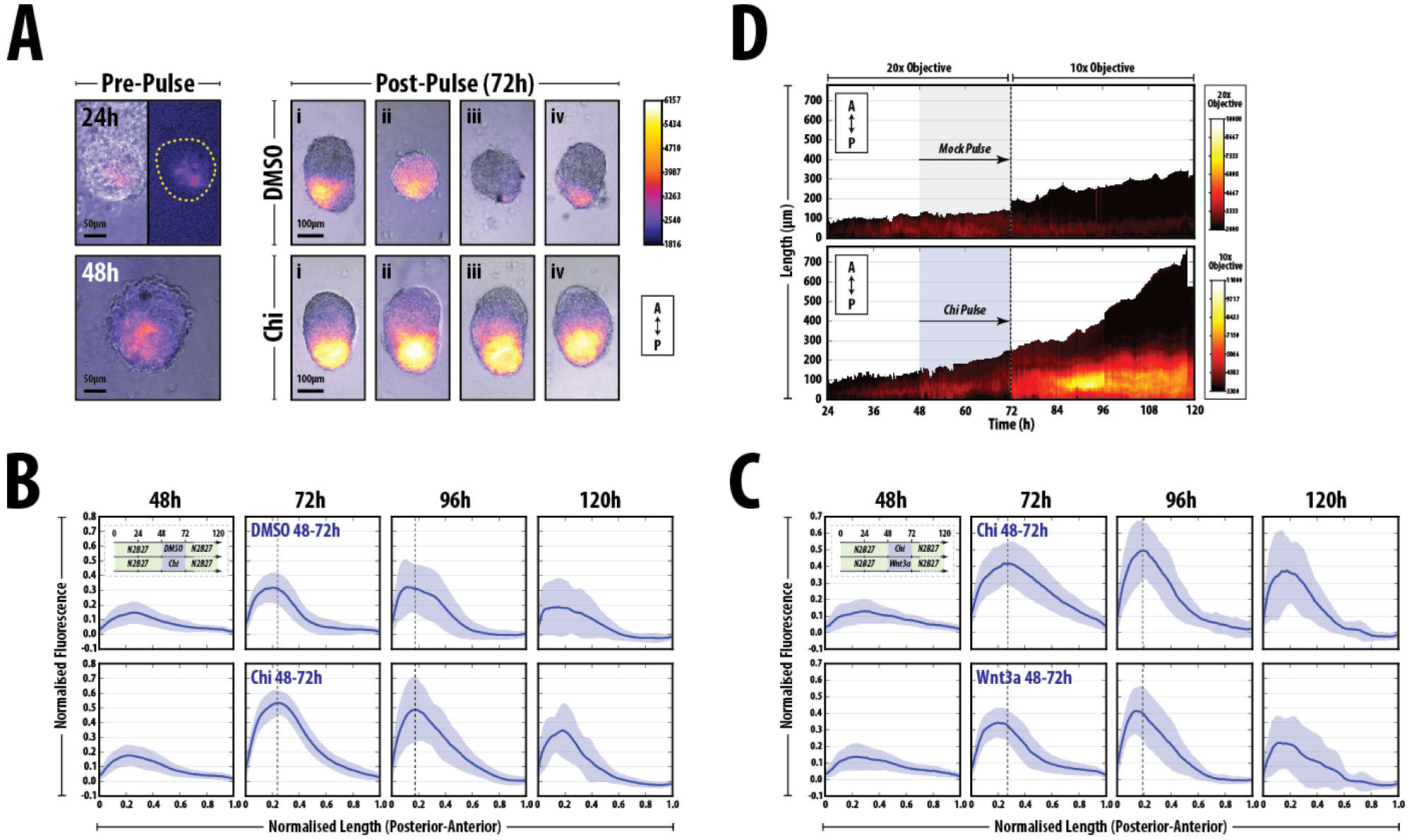
Symmetry-breaking and polarisation in the absence of signalling. *Gastruloids* generated from T/Bra::GFP were formed and maintained in N2B27 medium for the duration of the experiment and subjected to a 24h pulse of Chi or DMSO between 48 and 72h after aggregation (AA). (A) Morphology and expression of T/Bra::GFP at 24 and 48h prior to the Chi pulse (left), and examples (i-iv) of *Gastruloids* following either the Chi or DMSO pulse (right). Chi stimulation increases the robustness of the response and reproducibility of the phenotype. Anteroposterior orientation indicated (bottom right). (B) Quantification of T/Bra::GFP reporter expression in individual *Gastruloids* over time from one replicate experiment. The maximum length of each *Gastruloid* is rescaled to 1 unit and the fluorescence is normalised to the maximum fluorescence from the Chi condition. (C) Quantification of T/Bra::GFP *Gastruloids* following stimulation with either Chi or Wnt3a. (D) Live imaging of one representative *Gastruloid* subjected to a pulse of DMSO (top) or Chir (bottom) between 48 and 72h AA. Shown is the length of the *Gastruloid* over time (posterior = 0μm) and the fluorescence intensity of the reporter (colour). Early time-points (24-72h AA) imaged with a higher power objective. The schematic of the experimental design for *Gastruloid* stimulation is indicated as inserts in (B) and (C). Horizontal lines at 72 and 96h AA in (B) and (C) indicate the maximum fluorescence of the Chi pulse condition

Due to the reproducibility of these structures in culture and the ease with which they can be manipulated by the addition of specific signalling molecules, we were able to utilise this technique to assess how the initial patterning event in the *Gastruloids* is established.

Analysis of *Gastruloids* 24h AA revealed weak, spotty expression of T/Bra::GFP with a proportion already displaying signs of biased expression towards one pole (**Fig. 2**). By 48h AA in N2B27, the expression levels of T/Bra::GFP had risen and a more prominent polarisation and regionalisation of the reporter was observed in several *Gastruloids* (**Fig. 2A**). T/Bra::GFP was observed in one hemisphere of the *Gastruloids*, which tended to show a slight distortion in their shape from spherical to slightly ovoid as previously reported [13]. Addition of Chi at 48h increased both the levels of T/Bra::GFP and polarity by 72h AA (**Fig. 2**). If *Gastruloids* were treated with the vehicle control (DMSO), regionalised expression of the reporters could also observed, however the fluorescence levels were generally lower, with a higher degree of variation both within and between experimental replicates. This suggests that increased Wnt/*β*-Catenin signalling modulates the reproducibility and stability of the established pattern (Table S5).

To garner a better understanding of the heterogeneities in levels of T/Bra::GFP expression between the *Gastruloids* over time, we quantified the fluorescence in a posterior to anterior direction along the spine of the *Gastrulioids* (**Fig. 2B,C**; see Materials and Methods and [15]). T/Bra::GFP expression increased transiently over time, peaking at 96h AA (**Fig. 2B,C**), and was localised predominantly towards the posterior region (the origin) at all time-points (polarisation). Chi stimulation results in a lower standard deviation and a higher maximum fluorescence than the DMSO control and the expression of the reporter is maintained at the later time-points following Chi treatment (**Fig. 2B**). Stimulation Wnt3a resulted in a similar fluorescence trace compared with Chi (**Fig. 2C**), however the heterogeneity between individual *Gastruloids* is reduced and the higher expression is confined to a narrower region of the *Gastruloid* (increased polarity; **Fig. 2C**).

Next, to assess the evolution of T/Bra::GFP expression over time, we used live-cell microscopy to image the *Gastruloids* from 24 to 120h AA following a pulse of Chi or vehicle (**Fig. 2D**). Prior to stimulation, the size of the *Gastruloids* increased steadily over time with a concomitant increase in the levels of T/Bra::GFP to wards one pole, the expression of which became more apparent at approximately 36h AA (**Fig. 2D**). The addition of Chi at 48h did not have an immediate effect on the expression of the reporter until 72h AA, when a flash of T/Bra::GFP expression was observed throughout the aggregate; higher expression occurred towards the region that was already expressing T/Bra::GFP (**Fig. 2D**). Between 72-84h AA, T/Bra::GFP was down-regulated in the region that was previously not expressing the reporter and was up-regulated in the region that was already expressing T/Bra::GFP. This suggests that once a T/Bra::GFP domain is specified, only that region can stably respond.

Altogether, these data suggest that by 48h AA, the *Gastruloids* have developed an initial patterning, intrinsically driven, prior to the addition of signalling factors. Furthermore, increased Wnt/*β*-Catenin signalling by the addition of Chi increases the reproducibility both intra-and inter-experimentally, and robustness of the final phenotype and the patterning.

### 3.2 Gene Expression Analysis of Early Stage *Gastruloids*

The observation that the patterning of *Gastruloids* occurs in the absence of an external input led us to search for the expression of elements of the signalling pathways known to participate in symmetry-breaking in the embryo (**Fig. 1**). At 48h AA, *Gastruloids* express low levels of *FGF4, 5, Axin2, Wnt3, Nodal* and *Lefty1* (**Fig. 3**), all of which are expressed in the epiblast of the E6.0 embryo. Additionally, we detected no expression of *BMP4, Dkk* or *Chordin* with very low expression of *Cerberus* (**Fig. 3**). *Noggin*, an antagonist of BMP signalling, is not expressed in the early stages of development and its absence of expression serves as a baseline for the others (data not shown). This pattern is consolidated in the absence of any external input with increases in expression of *Nodal, Lefty1* and *FGF5*, decreases in *FGF4* expression and the emergence, at low levels, of *Wnt3a* at 72h AA (**Fig. 3**; 72h(D)). Exposure to Chi during 48 and 72 h AA leads to an increase in *Axin2, Nodal, Lefty, Wnt3a* and *Dkk*, all targets of Wnt signalling (**Fig. 3**; 72h(C)).

**Figure 3:**
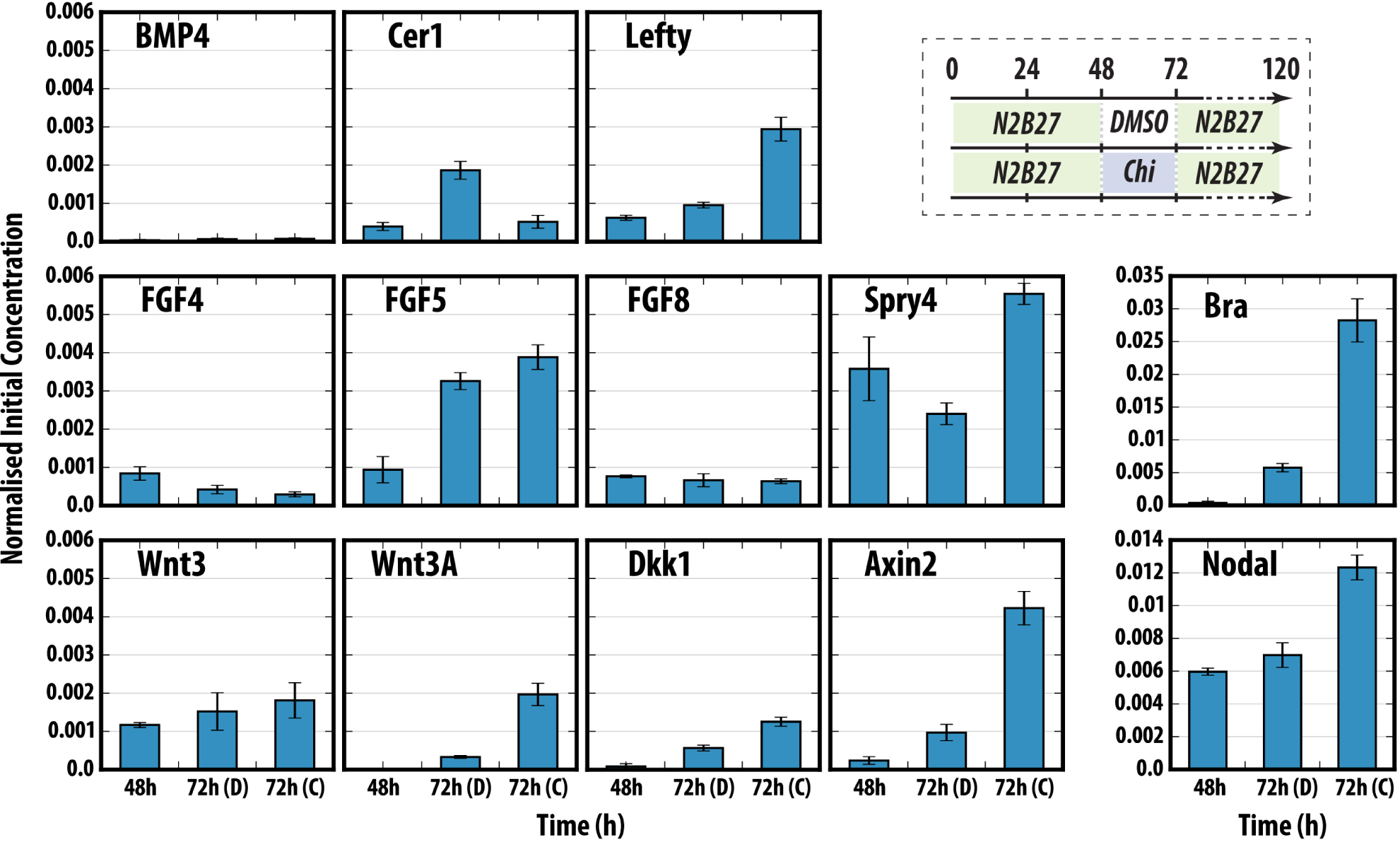
Quantitative PCR analysis of specific genes in early *Gastruloids* development, following a Chi pulse. *Gastruloids* generated from T/Bra::GFP mouse ESCs were harvested at 48 and 72h with (72h (C)) or without (72h (D)) a Chir pulse prior to RNA extraction and qRT-PCR analysis for the indicated genes. Data normalised to the housekeeping gene *Ppia* and is representative of 2 replicate experiments; error bars represent standard deviation of triplicate samples. Schematic of experimental design shown top right.

These results confirm that *Gastruloids* resemble the cells in the epiblast and also provide gene expression landmarks to correlate their development with events in embryos e.g. *FGF5* is a marker of the E5-E6.5 epiblast and *Wnt3a* and *Dkk* in the embryo, characterise the onset of gastrulation. Altogether these observations lead us to suggest that *Gastruloids* at 48h AA correspond, approximately, to E6.0(±0.5) in the embryo and 72h AA to E7.0 (±0.5). Further observations below confirm and extend this correspondence (**Fig. 3**).

### 3.3 Early Nodal expression and signalling are associated with the symmetry breaking event

The characterisation of the patterning and gene expression of early *Gastruloids* suggests that the Nodal/TGF*β* signalling pathway may be involved in the initial patterning event. To address this further, we assessed the pattern of expression of two reporters for Nodal activity in the *Gastruloids* from the early stages of aggregation (24h) up to 120h AA. *Gastruloids* were generated from ESCs containing a Nodal^condHBE::Y F P^ transcriptional reporter [17] (hereon referred to as the *Nodal Reporter*; (**Fig. 4**) or a Smad2/3 Nodal signalling reporter (AR8::mCherry) [18] (hereon referred to as the *Signalling Reporter* **Fig. 5**).

**Figure 4:**
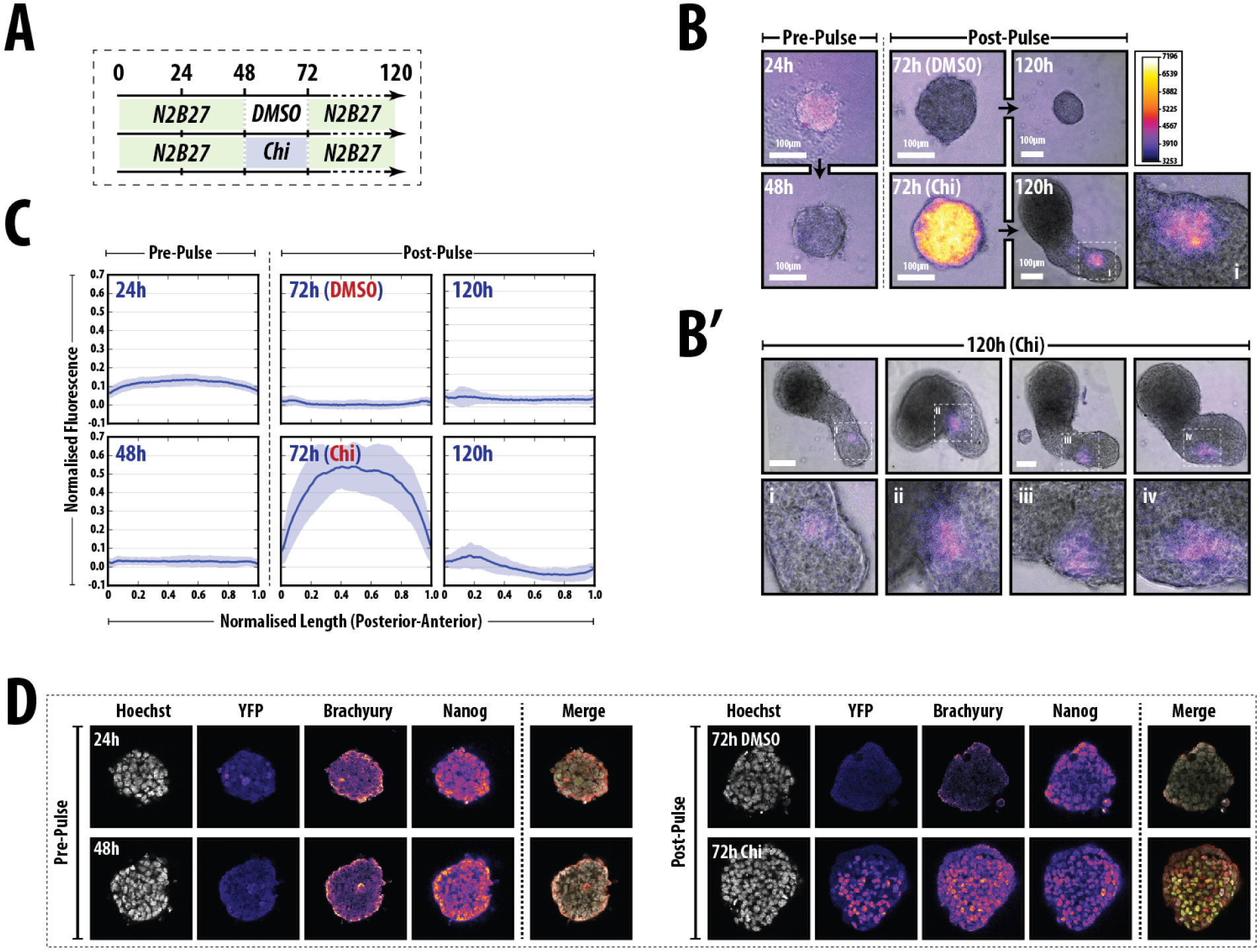
Expression of Nodal during early aggregate formation. (A) Schematic of the experimental design for *Gastruloid* stimulation. (B) *Gastruloids* formed from Nodal^condHBE::Y F P^ (Nodal reporter) ESCs either maintained in N2B27 or stimulated with a pulse of Chi or vehicle between 48 and 72h AA were imaged by wide-field fluorescence microscopy at 24, 48 and 72h AA. (B’) *Gastruloids* at 120h AA demonstrating a single region of Nodal expression slightly towards one side of the elongate; magnified regions (i-iv) indicated by hashed white box. (C) Quantitative analysis of the *Gastruloids* at the aforementioned time-points with their maximum length set to 1 unit and normalising their fluorescence to the maximum fluorescence of the Chi pulse at 72h AA. (D) A single slice through a confocal stack of the Nodal reporter *Gastruloids* stained for YFP, T/Bra and Nanog. See (**Fig. S2**) for the quantification of the confocal images. Notice the increase in Nodal expression after Chi stimulation.

**Figure 5:**
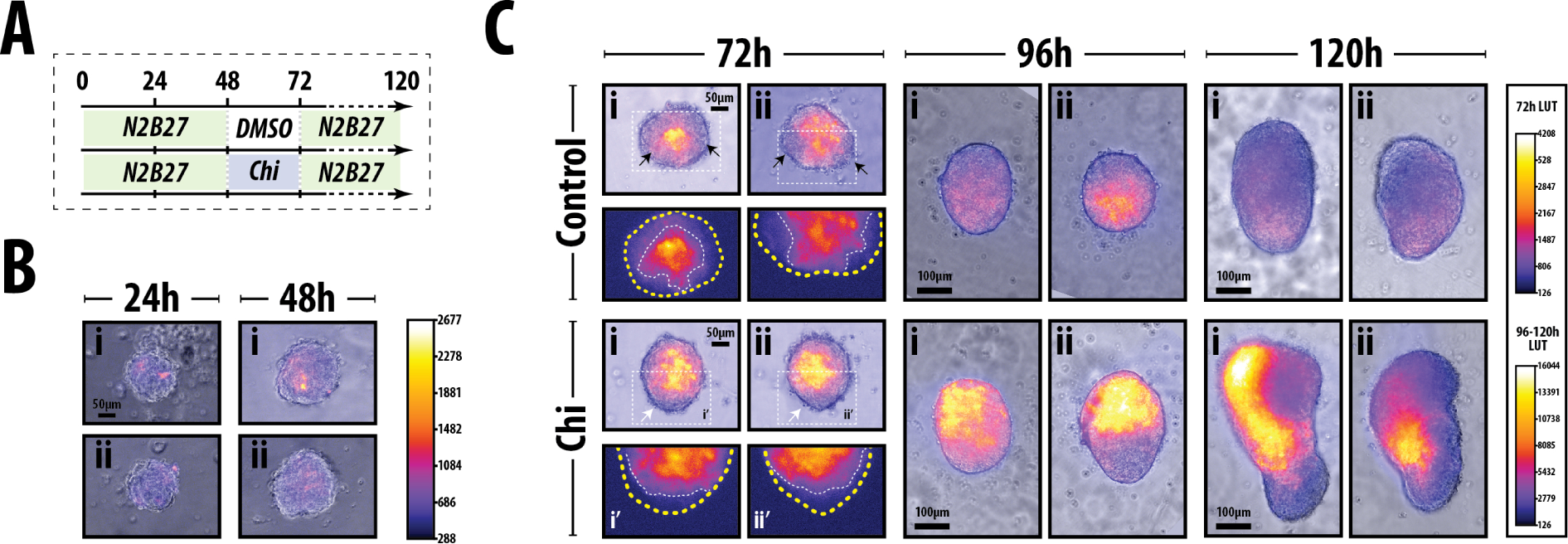
Expression of AR8::mCherry following Wnt/*β*-Catenin activation. (A) Schematic of the experimental conditions. (B) Expression of the AR8::mCherry reporter after 24 and 48h. (C) Expression of the reporter in two representative *Gastruloids* following either DMSO or Chi at 72, 96 and 120h. Magnified regions (hashed white line) at 72h are shown, indicating regions of the gastruloid that do not express the reproter. The 24h time-point in (B) was taken from a seperate experiment with identical imaging conditions as the 48h condition and all time-points in (C). The lookup tables for 24/48h, 72h and 96/120h has been rescaled to aid in the visualisation and localisation of the reporter when expressed at low levels.

At 24h AA, most of the cells within each *Gastruloid* expressed low levels of the Nodal reporter heterogeneously, with no evidence of a bias to any particular region and by 48h AA it was down-regulated throughout each *Gastruloid* (**Fig. 4A-C**). *Gastruloids* formed from the signalling reporter showed expression prior to the addition of Chi (**Fig. 5**) however, by contrast to the Nodal reporter, the signalling reporter was expressed in discrete punctate patterns that varied from *Gastruloid* to *Gastruloid* that increased, in number and intensity, between 24 and 48h AA.

Whereas continued culture in N2B27 maintained low expression of the Nodal reporter (**Fig. 4A,B**), following a Chi pulse between 48 and 72h AA, a dramatic increase in fluorescence throughout the whole *Gastruloid* was observed (**Fig. 4A,B**), that correlated with the expression within the epiblast at E4.5 and 5.5 [17]. Similarly, the signalling reporter was up-regulated following addition of Chi (**Fig. 5C**), however, at the end of the Chi pulse (72h AA), we observed in many *Gastruloids* a crescent free of the expression of the signalling reporter at the future posterior end (**Fig. 5C**). This pattern was also observed in the DMSO treated Gastruloids control, but regions of non-expression were more likely to appear in multiple locations (**Fig. 5C**). As with the Nodal reporter, vehicle treatment failed to up-regulate the signalling reporter to the same extent as Chi treated *Gastruloids* (**Fig. 5C**), indicating that Wnt/*β*-Catenin signalling was necessary for the enhancement and maintenance of Nodal signalling within one, polar region of the *Gastruloids*.

Between 96 and 120h AA, Nodal reporter *Gastruloids* treated with Chi confined Nodal expression to the elongating region and gradually reduced their expression (**Fig. 4B**,B’). By 120h AA these *Gastruloids* developed a polarised elongation and we note the emergence of high reporter activity in a group of cells centred at the anterior edge of the elongate; these cells may correspond to an attempt to establish a node as they mimic Nodal expression in the embryo [33, 34].

In support of this, in some instances, we observe low levels of the reporter restricted to one side of the aggregate, much as is the case for Nodal in the embryo (**Fig. 4B’i-iv**).

Finally, we sought to directly probe for the expression of T/Bra and Nanog within Nodal reporter *Gastruloids* to assess a) the correlation between Nodal expression and the acquisition of a posterior fate and b) to understand the timing of the initial symmetry-breaking event with respect to the loss of pluripotency (Nanog) and initiation of differentiation (T/Bra). Nodal reporter *Gastruloids* were fixed and stained for GFP, Brachyury and Nanog, and their expression assessed by confocal microscopy (**Fig. 4C**).

By 24h AA, Nodal was expressed at low levels in a heterogeneous manner, similar to that seen by wide-field microscopy (**Fig. 4D**, **Fig. S1**). Nanog was also expressed in a heterogeneous manner, with a broad distribution in fluorescence values (**Fig. 4D**, **Fig. S1**); there was no detectable above-baseline expression of Brachyury at this stage.

In *Gastruloids* maintained in N2B27 from 48h to 72h, Nodal expression remained low with no increase in the expression of Brachyury and a gradual reduction in the expression of Nanog (**Fig. 4D**, **Fig. S1**). This was reflected in the increasing correlation between Nanog, Brachyury and Nanog as their expression gradually reduced to background (Table S6). Stimulation with Chi from 48-72h increased the fluorescence of Nodal, Brachyury and Nanog in a heterogeneous manner, suggesting a commitment to a differentiated state (**Fig. 4D**, **Fig. S1**); the heterogeneous expression of Nodal in this region bears similarity to that in the embryo at E6.5[17]. Furthermore, there is evidence for expression of Nanog in this region at this time.

This suggests that events in the *Gastruloids* mimic events in embryos and that the period around 48-72h is related to the onset of gastrulation, and we would suggest here that it might encompass the E6.5-E7.5 in the embryo. In these conditions, the expression of Nodal and Brachyury was strongly correlated (Table S6).

### 3.4 BMP4 signalling is undetectable during Gastruloid Progression

In the embryo, genetic experiments have demonstrated the need for BMP signalling from the Extra embryonic regions for the expression and localisation of Nodal and Wnt signalling to the proximal posterior region of the embryo, however our data suggests that there is no extraembryonic component in the *Gastruloids*. To test this further, we investigated the activity of BMP signalling using a Smad1,5,7 reporter (IBRE4-TA-Cerulean; hereon known as the *BMP reporter*) [18] and measured its expression within *Gastruloids* treated either with DMSO or with Chi (48-72h AA) that had either been pre-treated with BMP4 or vehicle (**Fig. 6**).

**Figure 6:**
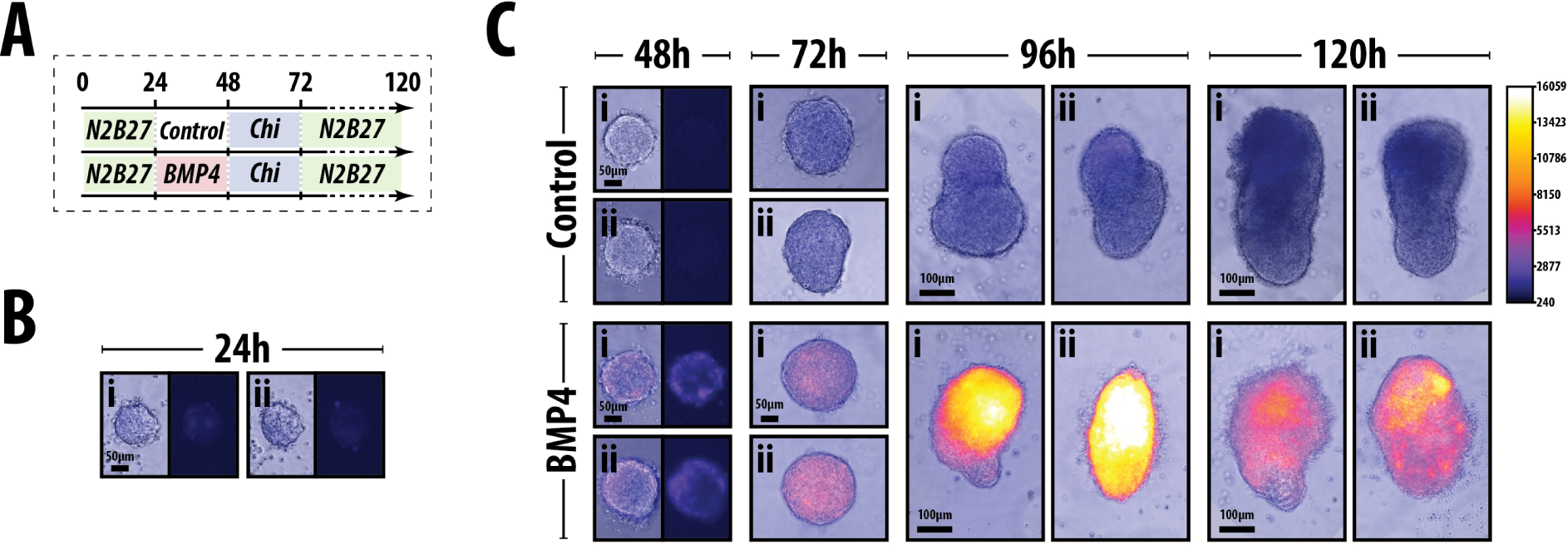
Expression of IBRE4-TA-Cerulean in Chi pulsed *Gastruloids* either pre-treated with vehicle or BMP4 for 24h. (A) Schematic of treatment and conditions. Cont; Control (HCl+0.1%BSA). *Gastruloids* imaged at 24h (B) and 47-120h (C) in the conditions described in (A). By 24h, there is very low BMP signalling. Following BMP pretreatment greatly increases the fluorescence of the reporter. Scalebar indicates 100 μm, lookup table displayed in (C) corresponds all images in this figure.

In *Gastruloids* only treated with a Chi pulse (control) we did not detect any significant increase in the expression of the BMP reporter from 24-120h AA. To test whether the cells are responsive to BMP at this stage, *Gastruloids* were pre-treated with BMP4 prior to the Chi pulse. BMP4 Pre-treatment greatly enhanced the activity of the reporter, with the majority of the expression located in the ‘anterior’ region (**Fig. 6C**).

Although it is possible that the reporter may not be sensitive enough to detect very low levels of signalling, these results demonstrate the lack of signalling in the *Gastruloids* and further support the notion that *Gastruloids* are made up of epiblast cells. In addition it also suggest that BMP signalling is not needed for the symmetry-breaking event itself nor the expression of T/Bra (see next section). Whatever low level of BMP signalling exists within the *Gastruloids*, it is not modified by Wnt signalling.

### 3.5 Functional analysis of signalling in the early patterning of *Gastruloids*

Our data suggest that interactions between the TGF*β*, FGF and Wnt/*β*-Catenin signalling before the pulse of Chi and as the *Gastruloids* polarise, mediate the symmetry-breaking event. To test this, we used agonists and antagonists of these pathways in the form of small molecules and recombinant proteins that target Nodal, BMP, FGF and Wnt/*β*-Catenin signalling on the day before the Chi pulse (**Figs. 7A**, **8A**, **9A**). It is to be noted that gentle mechanical and volumetric disruptions on the second day (24-48h AA) did not unduly affect the patterning and development of the *Gastruloids* (see Materials and Methods). As a read-out of the patterning events, we used T/Bra::GFP *Gastruloids* and recorded the fluorescence and morphology by widefield microscopy (**Figs. 7B**, **8B**, **9B**).

**Figure 7:**
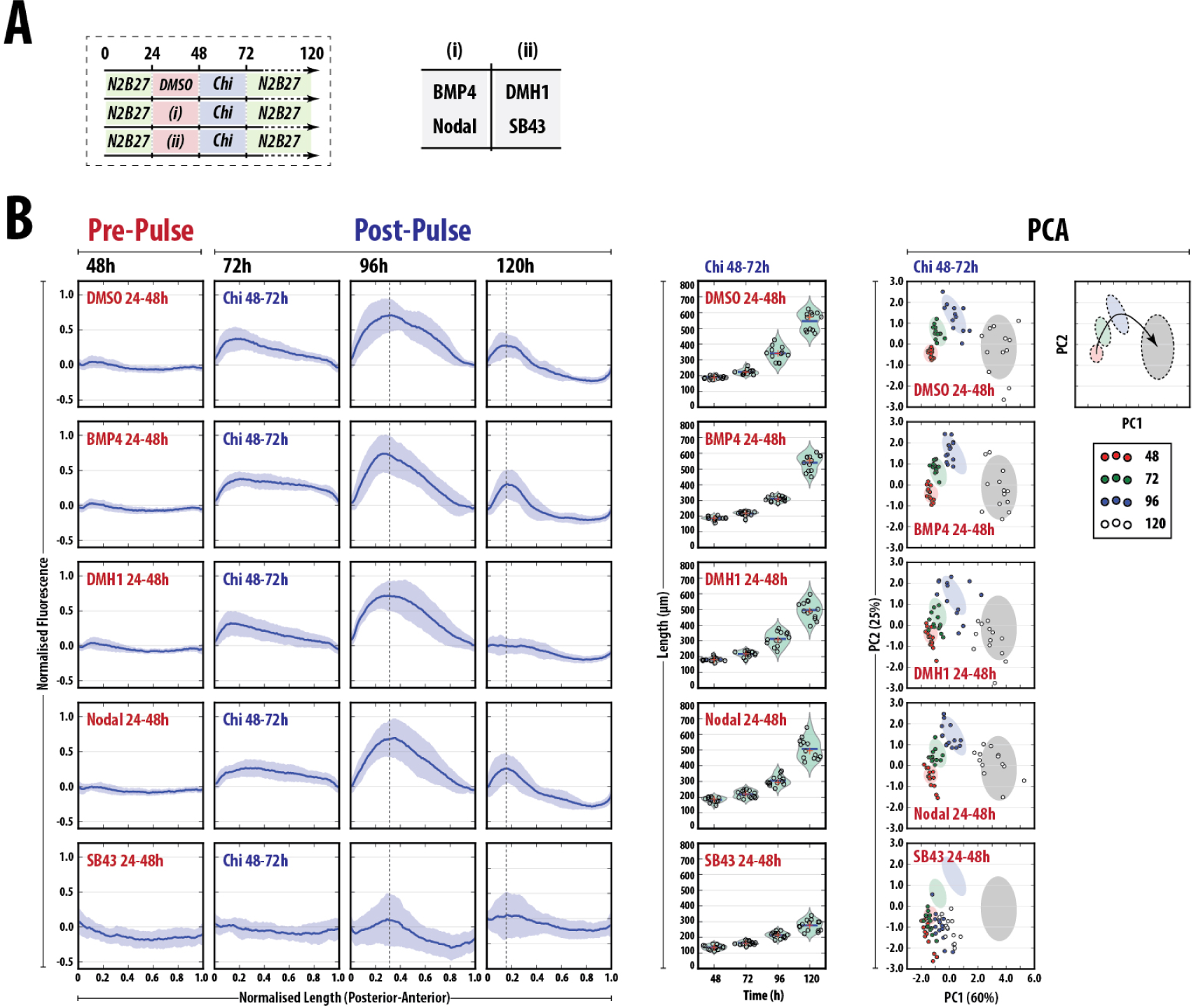
Quantitative analysis of T/Bra::GFP *Gastruloids* reveals importance of early Nodal signalling for correct patterning. (A) A schematic of the experimental design for *Gastruloid* stimulation. *Gastruloids*, aggregated in N2B27 (green shading) were treated with either vehicle, activators (i) or inhibitors (ii) of the Nodal and BMP signalling pathways shown in the table on the right, for 24h. Following wash-off of these factors, all *Gastruloids* were treated with a 24h pulse of Chi followed by sustained culture in N2B27. (B) Individual *Gastruloids* treated with vehicle, BMP4, DMH1, Nodal or SB43 were imaged by wide-field fluorescence microscopy and quantitatively analysed for the fluorescence (left panel) and length (middle panel). Principal Component Analysis (right) was used to reduce the dimensions of the data, revealing distinct clusters which corresponded to the specific time-points. The indicated percentage for the explained variance is denoted in the axis labels for PC1 and PC2. The fluorescence plots of the left of (B) were normalised to the maximum fluorescence of the control, and the maximum length of each *Gastruloid* was rescaled to 1 unit. Horizontal dashed lines at 72 and 96h AA in B indicate the maximum fluorescence of the Chi pulse condition.

**Figure 8:**
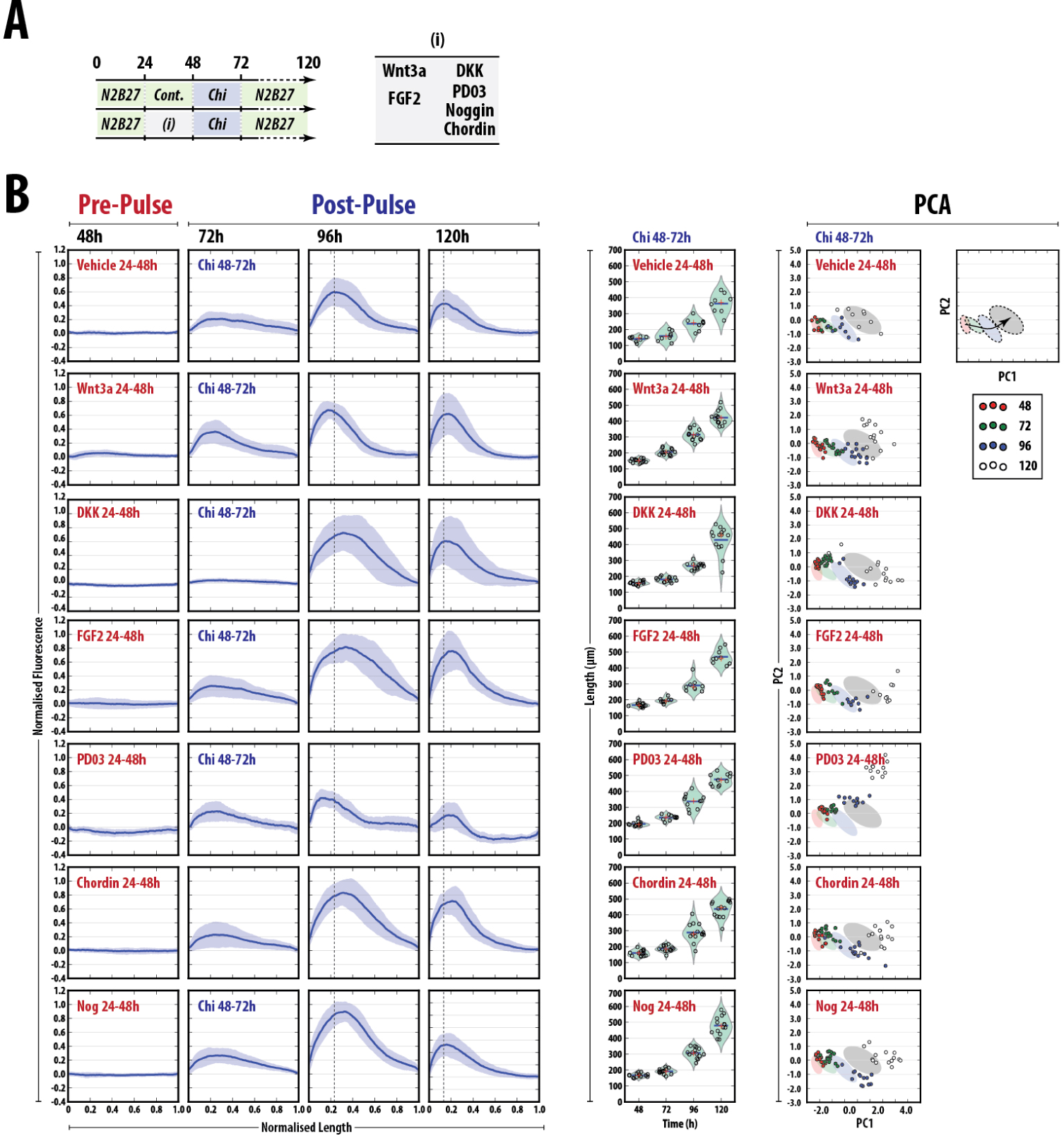
Effect of modulating Wnt, FGF and Nodal/BMP signalling prior to the Chi pulse. (A) Schematic representation of the experimental design (see **Fig. 7** legend). (B,C) Individual *Gastruloids* treated as labelled were imaged by wide-field fluorescence microscopy and quantitatively analysed for the fluorescence (left panel), length (middle panel), and their Area, circularity, perimeter and roundness measured (not shown). Principal Component Analysis (right) was used to reduce the dimensions of the data, revealing distinct clusters which corresponded to the specific time-points. The fluorescence plots of the left of (B) and (C) were normalised to the maximum fluorescence of the control, and the maximum length of each *Gastruloid* was rescaled to 1 unit. Horizontal dashed lines at 72 and 96h AA in B indicate the maximum fluorescence of the Chi pulse condition.

**Figure 9:**
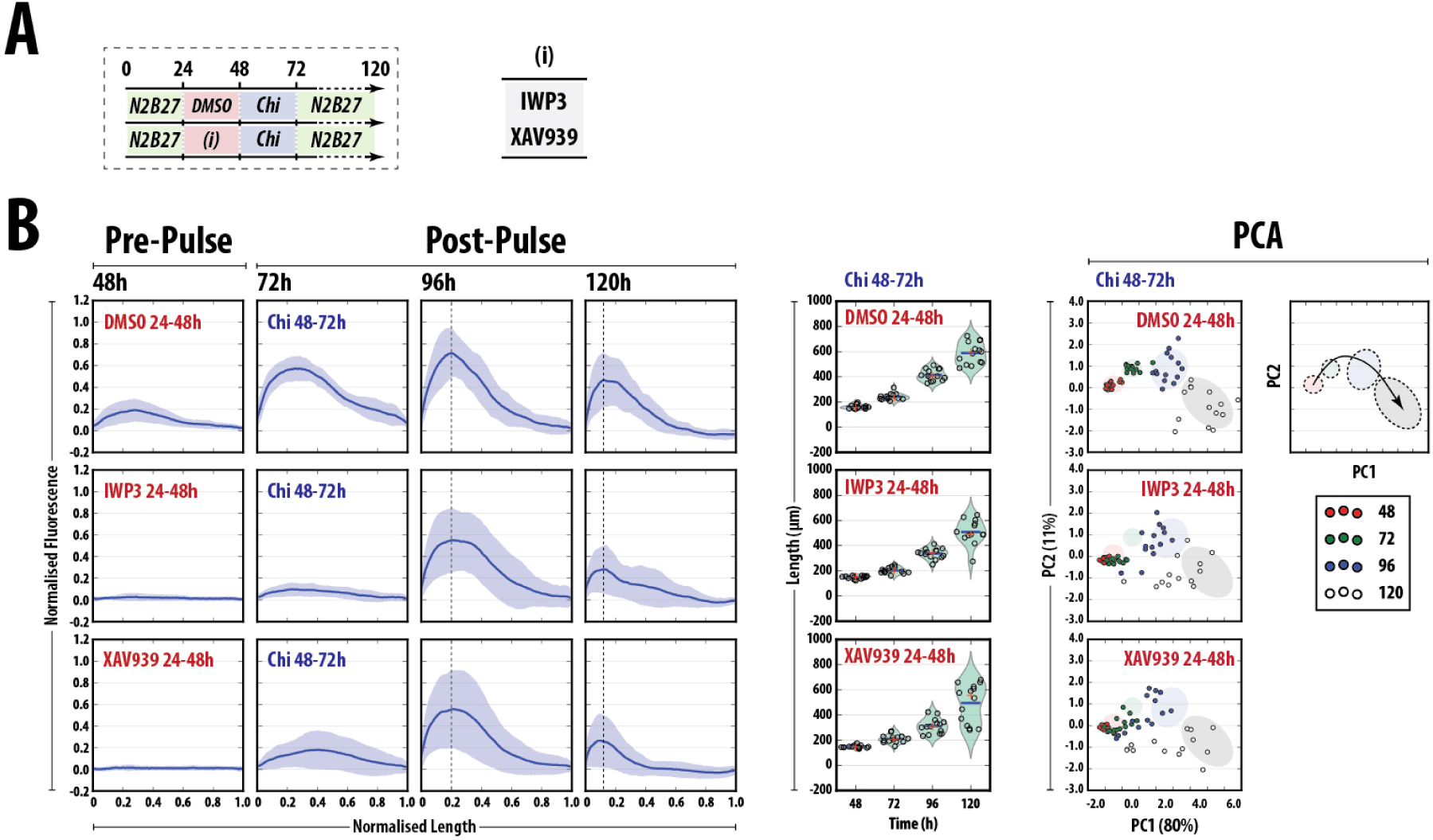
Quantitative analysis of T/Bra::GFP *Gastruloids* reveals defective pattern formation upon Wnt/*β*-Catenin disruption between 24 and 48h. (A) A schematic of the experimental design for *Gastruloid* stimulation. *Gastruloids*, aggregated in N2B27 (green shading) were pulsed with either DMSO, the Porcupine inhibitor IWP3 or the Tankyrase inhibitor XAV939 between 24 and 48h AA. Following wash-off of these factors (see materials and methods), all *Gastruloids* were treated with a pulse of Chi (48-72h AA) followed by sustained culture in N2B27. *Gastruloids* were imaged by wide-field fluorescence microscopy and quantitatively analysed for the fluorescence (B, left panel), length (B, middle panel) and their Area, circularity, perimeter and roundness. Principal Component Analysis (B, right panel) revealed that the discrete clusters that corresponded to the time-point in DMSO conditions were heavily disrupted following IWP3 or XAV939 pre-treatment. The fluorescence in (B) was normalised to the maximum fluorescence of the DMSO control. In addition, the length of each *Gastruloid* was rescaled so the maximum length had a value of 1 unit. Horizontal dashed lines at 72 and 96h AA in B indicate the maximum fluorescence of the Chi pulse condition.

As well as plotting the normalised fluorescence and the length of the Gastruloids for each specific treatment (left and middle panels of **Figs. 2A-C**, **7B**, **8B**, **9B**), we performed Principal Component Analysis (PCA) using the variables fluorescence (area under the curve), length, area, roundness and perimeter of the Gastruloids within each experimental replicate (right panels of **Figs. 7B**, **8B**, **9B**). By comparing the degree of clustering between individual *Gastruloids* and their trajectories through the PCA plot, we were able to discern the subtle and major differences each experimental treatment had upon the patterning and morphology of the *Gastruloids* compared with the internal control (i.e. 48-72h Chi pulse). Each experimental replicate included this internal control for quality control and to provide a matched baseline for morphology and *Gastruloid* progression.

In general, PCA of *Gastruloids* in control conditions revealed distinct clusters which corresponded to the time-point and the trajectories of these clusters through the plot were reproducible throughout replicate experiments (**Figs. 7B**, **8B**, **9B**, right panels). *Gastruloids* from early time-points were found to cluster tightly together (48h AA; red) with the final time-point (120h; white) being the most disperse; this may reflect heterogeneity between individual *Gastruloids* as they reach the end of their ability to be cultured for longer durations due to their increasing mass and propensity to adhere to the bottom of the wells at later times [15].

In all experiments (**Figs. 2A-C**, **7B**, **8B**, **9B**) and in agreement with previous observations [13], at 48h AA prior to the Chi pulse, a slight polarisation in the assumed-posterior region of the *Gastruloids* was observed which transiently increased in expression over time, peaking at 96h AA (**Fig. 7B**). Importantly, this pattern of expression is highly reproducible and allows us to use their expression as a landmark for the effect of the treatment on the initial patterning event. Additionally, their reproducibility allows us to us to use the standard deviation within each experiment as a measure of the effect of the treatment. In the description of the data below, we refer to the treatment before the Chi pulse as *pre-treatment*.

### 3.6 Nodal and FGF signalling are required for the symmetry breaking event

#### BMP signalling

Pre-treatment with BMP4 did not alter the expression levels or localisation of T/Bra::GFP at 48h AA (**Fig. 7B**). However following the addition of Chi, BMP4 pre-treatment enhanced the expression of T/Bra::GFP throughout the whole *Gastruloid* compared to the control, although by 96h, this was rapidly down-regulated and the expression pattern was essentially identical to the control by 120h AA (**Fig. 7B**, left panel). Progression of Gastruloid development following DMH1 pre-treatment was essentially identical to the control, although by 120h, DMH1 pre-treated Gastruloids were unable to maintain the expression of T/Bra::GFP (**Fig. 7B**, left panel). Whereas BMP4 pre-treatment showed a similar PCA distribution pattern to the controls over time with a slight increase in the spread in the second principal component at 48h AA, DMH1 pre-treatment resulted in a less coherent grouping of individual *Gastruloids* as time progressed (**Fig. 7C**).

These data suggest that BMP signalling is not involved in the early patterning of the *Gastruloids*, although as shown by DMH1 pre-treatment, the presence of low levels at these early stages (**Fig. 6**) may impact the ability to sustain T/Bra::GFP expression at later time-points.

#### Nodal signalling

Nodal pre-treatment (**Fig. 7B**) resulted in a similar fluorescence profile over time to that of the controls in terms of the fluorescence expression profile and the standard deviations between individual *Gastruloids* with no enhancement of the reporter at 48h, although, similar to BMP4 pre-treatment, Nodal pre-treatment resulted in a global up-regulation of T/Bra::GFP at 72h (**Fig. 7B**, left panel). Inhibition of Nodal signalling with SB43 completely abolished the expression of T/Bra::GFP, where only a small proportion of *Gastruloids* showed any expression at 96-120h AA (**Fig. 7B**, left panel), as well as a compromised pattern of growth, resulting in *Gastruloids* with a shorter final length. PCA revealed that although Nodal pre-treatment did not have a great effect on the overall trajectory of time-point clusters through the plot, there was a increase in the spread of the data points in both PC1 and PC2 at 48h AA (**Fig. 7B**, right panel). Inhibition of Nodal signalling with SB43 pre-treatment resulted in a complete disruption in the trajectories within the PCA, reflecting the observations with the fluorescence analysis (**Fig. 7B,C**).

As the *Gastruloids* show such a distinct phenotype following Nodal inhibition prior to the Chi pulse, these data reveal the absolute requirement for Nodal signalling in the initial patterning of *Gastruloids*.

#### FGF/MAPK signalling

To assess the role of FGF signalling, we pre-treated *Gastruloids* with either recombinant FGF2 or PD03 (MEK1/2 inhibitor) prior to the Chi pulse (**Fig. 8A**). Pre-treatment with FGF2 resulted in a T/Bra::GFP fluorescence profile with slightly increased heterogeneity between individual *Gastruloids* at 48h AA compared with the vehicle control (**Fig. 8B**, left panel). By 72h AA, the expression of T/Bra::GFP following FGF2 pre-treatment was essentially identical to the vehicle control, however, at 96h, FGF pre-treatment resulted in a higher maximum fluorescence than the control with the pole of expression less well defined (i.e. the expression was spread over a larger posterior region; **Fig. 8B**, left panel). This increased expression levels of the T/Bra::GFP reporter was maintained at 120h AA and although the region of expression had become more localised to the posterior region, the expression levels were augmented with respect to the controls (**Fig. 8B**, left panel). Interestingly, although Inhibition of FGF signalling by PD03 pre-treatment had little effect on the expression of T/Bra::GFP within the *Gastruloids* between 48 and 72h, *Gastruloids* at later time-points (96-120h AA) were severely retarded in their ability to up-regulate the reporter to the same extent as either the control or FGF2 pre-treatment condition (**Fig. 8B**, left panel). In both pre-treatment conditions, the gradual increase in the length of the T/Bra::GFP *Gastruloids* was not significantly altered over time when compared to the control(**Fig. 8B**, left panel).

In addition to the T/Bra::GFP cell line, we utilised an FGF reporter (Sprty4::H2B-Venus) mouse ESC line to assess FGF/MAPK signalling and assessed its expression following pre-treatment with either FGF2 or PD03 (**Fig. S2**). Vehicle treatment resulted in a gradual increase in the FGF reporter over time with a slight bias towards the posterior region at 48 and 72h AA (**Fig. S2**), with a burst of expression throughout the whole *Gastruloids* at 96h AA. Interestingly, FGF2 pre-treatment did not have an effect on the average amplitude of the fluorescence between 48 and 72h AA, although posterior pole of expression was more defined than the control, and treatment with PD03 resulted in a gradual decline of reporter expression over time (**Fig. S2**). By 96h *Gastruloids* pre-treated with FGF2 resulted in much higher levels at 96h than the control. Interestingly, the large increase and decrease in expression levels of the reporter following FGF2 or PD03 pre-treatment respectively occurred at the same time as the broadening and reduction (respectively) of the T/Bra::GFP. These data suggest that although FGF signalling does not impact the initial patterning of the *Gastruloids*, it is involved in the maintenance of T/Bra::GFP expression at later stages.

Analysis of the *Gastruloids* using PCA revealed FGF2 pre-treatment to be broadly similar to the controls, with discrete clusters corresponding to the time interval following the same trajectory set out by the controls (**Fig. 8B**, right panel). However, FGF2 pre-treatment resulted in a more rapid progression, where earlier time-points (e.g. 72h AA) occupied the regions reserved for the 96h time-point (as defined by the controls; **Fig. 8B**, right panel). In addition, the heterogeneity between *Gastruloids* was greatly reduced and the clustering appeared tighter than the controls (**Fig. 8B**, right panel). Conversely, PD03 pre-treatment greatly disrupted both the usual progression, indicating the requirement of FGF2 for *correct* passage through the plot (**Fig. 8B**, right panel).

Taken together, these results indicate that FGF signalling has limited effect on the either initial patterning of the *Gastruloid*, the up-regulation of T/Bra::GFP fluorescence or progression of the *Gastruloids*, however the low, endogenous levels that are present at these early time-points are essential for constraining the domain of expression at the posterior region and the maintenance of increased expression at later time-points. It may be the case that FGF signalling is acting to lower the threshold of the response.

#### Wnt/*β*-Catenin signalling

To assess the effect of Wnt/*β*-Catenin signalling, we pre-treated *Gastruloids* with the recombinant proteins Wnt3a or DKK1, (**Fig. 8**) or small molecule inhibitors (IWP3: to inhibit the secretion of all Wnts [35]; XAV939; to increase *β*-Catenin degradation through tankyrase inhibition[36]; **Fig. 9**).

Treatment with Wnt3a resulted in a slightly enhanced posterior activation of T/Bra::GFP at 48h AA compared with the vehicle control (0.1% BSA in PBS) which continued into the 72h time-point (**Fig. 8B**). As time progressed, the Wnt3a pre-treated *Gastruloids* showed a narrower region of T/Bra::GFP expression, with the peak expression occurring more posteriorwise than the vehicle control, displaying less variation between individual *Gastruloids* (**Fig. 8B**). Although peak T/Bra::GFP expression occurred at 96h AA following both vehicle and Wnt3a pre-treatment, Wnt3a tended to increase the fluorescence of the reporter to a greater extent than the control(**Fig. 8B**). Interestingly, whereas control *Gastruloids* generally reduced T/Bra::GFP expression by 120h, Wnt3a pre-treatment maintained the expression, and appeared to confine its expression to a narrower region within the posterior, albeit with a higher standard deviation(**Fig. 8B**).

Inhibition of LRP6-mediated Wnt signalling by pre-treatment with recombinant Dkk1 resulted in an identical expression profile of T/Bra::GFP to vehicle control at 48h, however following Chi stimulation the fluorescence activation usually observed at 72h AA following vehicle or Wnt3 pre-treatment was abolished in the Dkk1-pre-treated *Gastruloids* (**Fig. 8B**). By 96h, the expression of the reporter had recovered, displaying maximum fluorescence at similar levels to the vehicle and Wnt3a conditions, however, the heterogeneity between individual *Gastruloids* was greatly increased and the polarisation was less defined (i.e. the fluorescence was less concentrated at the posterior region; **Fig. 8B**). After 120h in culture, the fluorescence of Dkk1 pre-treated *Gastruloids* was broadly similar to the Wnt3a pre-treatment with a much higher degree of variation between individual *Gastruloids* than the vehicle control (**Fig. 8B**).

Pre-treatment with either IWP3 or XAV939 delayed the onset and magnitude of T/Bra::GFP expression in a similar manner to that of Dkk1 (**Fig. 9B**, left panel). In some replicate experiments in the XAV939-pre-treated *Gastruloids*, the fluorescence expression, although still generally located posterior-wise, displayed a greater degree of variation with respect to the DMSO control (data not shown). Similar to the Dkk1 pre-treated *Gastruloids*, by 96 and 120h AA, the pattern of T/Bra::GFP fluorescence was broadly similar to the controls although a greater deal of variation was observed between the *Gastruloids* pre-treated with XAV939 and IWP3 and a slightly lower average expression level (**Fig. 9B**, left panel). Interestingly, *Gastruloids* pre-treated with IWP3 (which inhibits all Wnt secretion) was unable to maintain the expression of the reporter at 120h compared with Dkk1 inhibition (inhibits at the level of LRP6), suggesting a requirement for non-canonical Wnts in the maintenance of T/Bra::GFP fluorescence.

Comparison of the lengths of the *Gastruloids* following either vehicle, Wnt3a or Dkk1 pre-treatment showed no differences in the progression of the average lengths, however Wnt3a pre-treatment reduced the heterogeneity between individual *Gastruloids* whereas Dkk1 pre-treatment resulted in a greater variation at 120h. However, there was a slight disruption in the maximum lengths obtained at 120h AA following IWP3 or XAV939 pre-treatment, with a slightly lower average length and a larger spread of final lengths (**Fig. 9B**, right); this was more pronounced with XAV939 pre-treatment.

PCA analysis of *Gastruloids* pre-treated with Wnt3a and Dkk1 revealed similar trajectories to their control (**Fig. 8B**, **9B**). Interestingly, *Gastruloids* pre-treated with Wnt3a from the 72 and 96h time-points were able to explore region usually reserved for the 96h and 120h time-points respectively (**Fig. 8B**, **9B**), suggesting that Wnt pre-treatment may enhance the response to Chi. A number of subtle differences between the methods of Wnt inhibition by Dkk1, IWP3 or XAV939 pre-treatment was also revealed by the PCA analysis (**Fig. 8B**, **9B**). Whereas pre-treatment with Dkk resulted in a similar trajectory to the control (although with a greater shift in the positive direction in PC1 at 72h AA), the trajectories of IWP3 and XAV939 pre-treated *Gastruloids* was greatly disrupted compared to the controls and the data-points which correspond to specific time intervals were less coherently clustered (**Fig. 9B**). Prior to Chi stimulation, the clustering of the 48h time-point of both IWP3 and XAV939 was dispersed along the PC2 to a greater extent than the control which was unable to be recovered after the Chi pulse (72h). By 96h, IPWP3 and XAV939 pre-treated aggregates had largely recovered in terms of their positioning relative to the control, however XAV939 had not fully recovered and data-points corresponding to individual *Gastruloids* were spread throughout PC1, occupying regions reserved for earlier time-points (**Fig. 9B**). After 120h in culture, neither IWP3 or XAV939 pre-treated *Gastruloids* were able to reach the same region of the PCA as the controls; this is reflected in the fluorescence traces which were unable to maintain T/Bra::GFP expression to the same extent as the control (**Fig. 9B**). These data suggest that the effects of IPW3 and XAV939 are greater than Dkk1-mediated Wnt inhibition, when many of the variables (length, fluorescence, roundness, area etc.) are taken into account. This further suggests a requirement of ‘non-canonical’ (i.e. *β*-Catenin-and LRP6-independent) Wnt signalling at later stages, the secretion of which was inhibited by IWP3. Upon closer inspection of the expression of the T/Bra::GFP reporter within the Gastruloids, we found that whereas T/Bra::GFP expression is maintained at the elongating tip of the control *Gastruloid* (as previously described), treatment with IWP3 resulted the expression of the reporter along a mid-line region of a small subset of *Gastruloids* per experimental replicate (**Fig. S3**, indicated by white arrows); this phenotype was less pronounced in XAV939 pre-treatment conditions. Only one *Gastruloid* with Dkk pre-treatment showed this phenotype (data not shown).

#### Section summary

Although there was slight variation between replicates (in terms of expression levels and the internal standard deviations), these observations indicate that whereas BMP and FGF signalling has minimal effect at the onset of the initial patterning event, Nodal signalling is absolutely essential to allow the not only the expression of T/Bra::GFP but possibly its correct placement within the aggregate and Wnt/*β*-Catenin signalling is required in this early period (24-48h AA) for the correct timing of expression. The requirement of BMP signalling later in the *Gastruloid* development suggests two fundamental stages of *Gastruloid* development: one base on establishing the pattern which is Nodal and Wnt dependent and the second on its growth, and elongation involving a late requirement of both FGF2 and BMP signalling, and ‘non-canonical’ (e.g. LRP6-independent) Wnt signalling. In addition, these results clearly demonstrate the reproducibility of the *Gastruloid* phenotype not only within experiments but throughout replicate experiments; this measure of bother intra-and inter-experimental reproducibility has not to our knowledge been demonstrated other organoid culture techniques [37].

## 4 Discussion

In the mouse, the anteroposterior axis is established early in the cup-shaped zygote through the localisation of Nodal, BMP and Wnt signalling to a proximoposterior region of the embryo that becomes the posterior pole, where gastrulation is initiated as reflected in the expression of T/Bra (**Fig. 1**). Here we have shown that *Gastruloids*, embryonic organoids derived from small aggregates of mouse ESCs, undergo symmetry-breaking and gene expression polarisation in a manner that mirrors events in embryos [13–15, 37]. However, in contrast with embryos that use a sequence of interactions between extraembryonic and embryonic tissues to pattern themselves, *Gastruloids* are composed only of embryonic tissue and reveal an intrinsic capacity for axial organisation that is driven by interactions between Nodal and Wnt signalling. Analysis of signalling reporters in the *Gastruloids* reveals clear Wnt/*β*-Catenin and Nodal/Smad2/3 activity from 24-48h AA which often becomes localised to one pole of the aggregate and correlates with the onset and localisation of *T/Bra* expression. Furthermore, as in the embryo, this pattern of *T/Bra* expression is dependent on Nodal and Wnt signalling: exposure of *Gastruloids* to inhibitors of these pathways between 24 and 48h AA abolishes or reduces these patterns.

The absence of extraembryonic tissues is reflected in the absence of expression of Trophoectoderm and Visceral Endoderm markers (*BMP4, Cerberus* or *Dkk1*) and the absence of BMP reporter activity (**Figs. 3**, **6**). Furthermore, addition of BMP to the culture, has little effect on the patterning process other than a transient increase in the levels of T/Bra through alterations of Nodal and Wnt signalling (data not shown). In the embryo, BMP is deployed in the Extraembryonic Ectoderm and regulates the expression of Nodal and Wnt3 in the proximal part of the embryo [38, 39]; a similar effect has been reported in differentiating ESCs [40, 41] and can account for the observed effect BMP on the *Gastruloids*.

The strict dependence of Nodal and Wnt expression on BMP signalling in the embryo and the requirement for their antagonists in the symmetry-breaking event, raise questions about the molecular origin of polarity in our experiments. In the embryo Nodal plays a key role in the establishment of AP polarity [42, 43] and this also appears to be the case in the *Gastruloids* where in the absence of Nodal signalling there is neither localisation of T/Bra::GFP expression nor clear polarisation of the *Gastruloid*. Furthermore, in the embryo successive patterns of *Nodal* expression outline the emergence of the AP axis: it is first expressed throughout the E4.5 blastocyst and the E5.5 epiblast, where it then becomes restricted, first to its proximo-posterior region and shortly afterwards, in a Wnt/*β*-Catenin signalling dependent manner, to the Primitive Streak [44]. The patterns of Nodal expression in the *Gastruloids* are reminiscent of these: an onset of low ubiquitous expression coincident with the extinction of Nanog expression, a Wnt/*β*-Catenin dependent rise between 48 and 72h AA and then a restriction to the elongating region (**Fig. 4**). These patterns of Nodal expression are associated with Smad2/3 activity (**Fig. 5**). Interestingly, around 96h AA we observe the emergence of a spot of Nodal expression at the anterior border of the elongating region which could correspond to a structure similar to the node in the embryo [45, 46]. This possibility is supported by the observation of Smad2/3 reporter activity between 96 and 120h AA on one side of the Gastruloid (see **Fig. 4**, **5**); in some instances it is possible to see a similar but faint pattern of Nodal expression (data not shown). This pattern of activity is reminiscent of the Left-Right asymmetry expression of Lefty and Nodal in the embryo [47] and suggests that the *Gastruloids* not only develop an AP axis but also exhibit bilateral asymmetry.

The self organised patterning of the *Gastruloids* is, at first sight, surprising but might reflect a situation, latent in the embryo, which generates asymmetries in gene expression within the E5.0 Epiblast, before the restriction of primitive streak initiation to the proximal posterior part of the embryo [48, 49]. This possibility is supported by some observations. For example, the epiblast of embryos double mutant for *Cerl* and *Lefty1* is not unpatterned and exhibits a residual asymmetry in *Nodal* expression and, sometimes, double axes [49, 50]; this suggests that the epiblast can pattern itself in the absence of the extraembryonic repressors. Similarly, loss of *Dkk1* and gain of function *β*-Catenin do not alter the axial organisation of the epiblast, though they eliminate the head [51]. It could be argued that there are functional redundancies in the antagonists of Wnt and Nodal signalling and that this explains the *partial polarity* in *Dkk* and *Cerb/lefty* double mutants. However, this does not account for the observed phenotypes and suggests that these antagonisms are not drivers of AP patterning.

Our results argue for an independent and intrinsic self-patterning ability in the epiblast, and we would like to suggest that the role of the visceral endoderm and the extraembryonic ectoderm (trophoectoderm) is not to break the initial symmetry but to bias and stabilise an intrinsic symmetry-breaking event in the epiblast. Crucially, to position the initiation of gastrulation precisely and reproducibly to one end of the proximal part of the conceptus: near the extraembryonic Ectoderm. The notion that within organisms, intrinsic patterning activities of individual structures are biased and contribute in these complex connected structures has been discussed in other related contexts [37]. The reason for this is likely to lie in the fact that the first cells to leave the Primitive Streak will allocate themselves within the Extraembryonic Ectodermal territory to give rise to the allantois and the chorion [52, 53], and that positioning the initiation of gastrulation to the proximal domain of the embryo facilitates these cells to find their target. A process solely determined by a spontaneous symmetry-breaking event would position this point indiscriminately and therefore not contribute efficiently to the general organisation and structure of the conceptus. Thus, it is possible that the interactions between embryonic and extraembryonic tissues during axial specification are related to the interactions between the embryo and the mother more than between the organism and the embryo.

### Nodal signalling and AP polarity

Our results suggest that symmetry-breaking in *Gastruloids* is reliant on Nodal that is expressed from the moment of aggregation. Nodal is known to play a critical role in the establishment of the AP axis in all vertebrates [54, 55] and, together with its transcriptional target and feedback inhibitor Lefty1, has been shown to have an intrinsic symmetry-breaking ability [56–58]. Our results suggest that it is this characteristic that probably mediates the symmetry-breaking in the *Gastruloids*. We observe low levels of Nodal signalling as soon as the aggregate forms and the expression of Nanog, a pluripotency marker, is extinguished. This has been observed in ESCs [59] and might reflect the switch between pluripotency and differentiation. Later, between 24 and 48h AA we detect a range of patterns of the Smad2/3 reporter, with frequent small patches of high levels of activity polarised to one side. We also detect low levels of *Lefty1* expression, which is a target of *Nodal* and its main partner of its self patterning capacity. Furthermore, *Nodal* is a target of Nodal signalling [57] and also has an input from Wnt signalling [60]; this is also recapitulated here where Wnt/*β*-Catenin signalling does lead to an increase in *Nodal* expression (**Fig. 4**). At 48h AA, we observe patches of low level T/Bra::GFP expression and Wnt/*β*-Catenin signalling. In the absence of any extrinsic signals, this pattern evolves and by 72h AA it is possible to observe T/Bra::GFP and Wnt/*β*-Catenin signalling localised to one pole and *Nodal* expression to the elongating region in a variable domain of the initial *Gastruloids*. In the embryo this pattern of expression marks the start of gastrulation at E6.2 and therefore, is consistent with our estimate that 48h AA is about E5.5 as this maps the interval between 48 and 72h AA to approximately between E5.5 and E6.5 (**Fig. 10**).

In the embryo, expression of Nodal at E5.5 appears heterogeneous (see Fig. 1A in [44]) as it is at 48h AA in our experiments. This pattern and its evolution might represent the development of a Reaction-Diffusion (RD) system that drives the symmetry-breaking event both, in the embryo and the *Gastruloid* [56, 57, 61]. The dependence of an intrinsic symmetry-breaking event on Nodal signalling in the embryo is supported by the observation that the patterning defects observed in *Cerberus, Lefty* double mutants are corrected by reducing the dosage of Nodal in the mutant embryos [49]. It may be that the interactions between the extraembryonic and embryonic tissues raise the threshold for Nodal activity in the embryo and that this is part of the proximodistal biasing mechanism that we have discussed. The notion that interactions between tissues change the thresholds of signalling events involved in pattern formation in order to positions structures relative to each other might be a principle that emerges from these and related studies [37].

In the embryo, Nodal signalling requires Furin convertase activity which, during E5.0 and E6.0 is provided by the extraembryonic ectoderm [60]. In principle this activity should be missing in the *Gastruloids* but, as they pattern themselves in a Nodal dependent manner and we observe Nodal dependent activity of a Smad2/3 reporter, there must be some residual convertase activity or, as has been suggested, the Nodal precursor is able to signal through Activin receptors [38]. Further work should clarify this.

### The role of Wnt signalling

The establishment of the AP axis in vertebrate embryos is marked by the onset of gastrulation (represented by the primitive streak in amniotes) at the posterior midline. Experiments in *Xenopus* revealed that, in addition to Nodal, Wnt/*β*-Catenin signalling plays a key role in this process [63–65], and similar requirement has been revealed in chicken embryos, where the initiation of gastrulation requires a synergy between Nodal and Wnt/*β*-Catenin signalling [66, 67]. Analysis of mouse mutants for elements of Wnt signalling supports this interaction [7, 8]: while gain of function of Wnt signalling can generate an ectopic axis [68, 69], loss of function leads to defects in polarity and axial extension [70]. Nodal and Wnt signalling synergise in the induction of *T/Bra* expression and the initiation of the primitive streak [66, 67] but, how interactions between these signalling pathways lead to a robust AP axis remains open to discussion. Our experiments provide some insights into the nature of these interactions.

At 48h AA, the patterning of *Gastruloids* appears to be a stochastic event. Whether a *Gastruloid* will elongate or not in the next 48 hours appears to correlate with the localisation of *β*-Catenin/TCF, Smad2/3 signalling and T/Bra expression to one pole of the *Gastruloid*, and varies from experiment to experiment. Surprisingly, exposure to Chi or Wnt3a for 24h between 48 and 72h AA leads to an almost uniform polarisation of both T/Bra::GFP and Wnt signalling as well as elongation with T/Bra::GFP at the tip of the *Gastruloid* across a population (**Fig. 2**, **7**). This observation is surprising as there is no polarised exposure to Chi or Wnt and yet, the response is localised and triggers a precisely spatially orientated event. Furthermore, time-lapse imaging of the behaviour of *Gastruloids* during and after exposure to Chi reveals a transient global response in the form of ubiquitous expression of T/Bra::GFP, which then relaxes to the posterior region of the *Gastruloids*, to where is was before the signalling pulse (**Fig. 2D**). It seems as if the only region able to respond would be the one that had been already chosen.

There are several examples in which ubiquitous tonic activation of Wnt signalling results in a localised response or, in many instances, is capable of rescuing a Wnt signalling mutants [71–76]. Furthermore, loss of Wnt signalling reduces but does not abolish the expression of genes under its control [70, 77] and in some instances, has no effect on the pattern [78, 79]. One explanation for these observations is that rather than working as a morphogen or directing the output of a gene regulatory network (GRN), Wnt signalling controls the signal to noise ratio of the processes it is associated with [75, 76]. In the case of the *Gastruloids*, this would be the Nodal-driven symmetry-breaking process and its effect on T/Bra which is a target of Wnt/*β*-Catenin signalling. In support of this, exposure to the Nodal/ALK4/7 signalling inhibitor SB43 during 24-48h AA abolishes the response to Chi and the polarisation of the aggregate. Furthermore, increases in Wnt/*β*-Catenin, but not Nodal, signalling will rescue losses of Wnt signalling during the same period. This suggests that Nodal is the main driver of the symmetry-breaking event during 24-48h AA and that Wnt/*β*-Catenin signalling modulates its effects.

Our results support the hypothesis that a key role of Wnt signalling in developmental events is to act as a filter in cell fate decisions that regulates the ratio of signal to noise of specific processes [75, 76, 80] and suggest that in the *Gastruloids*, the effect of Wnt/*β*-Catenin signalling is to stabilise the molecular and cellular events that have taken place during 24-48h AA, principally the activity of Nodal/Smad2/3 activity. Interestingly, application of Chi after disrupting Wnt signalling during this period results in a process with much more variability than the wild type situation (**Fig. 9**). One interpretation of this observation is that, in the absence of Wnt/*β*-Catenin signalling, the output of Nodal/Smad2/3 activity is noisy and this is what is amplified by Chi between 48 and 72h AA i.e. Wnt/*β*-Catenin signalling acts as an amplifier/filter of previous events rather than the driver of the symmetry-breaking, which is likely to be Nodal. This situation is supported by the phenotype of Wnt3 mutant embryos which exhibit initial, weak but localised expression of T/Bra that then progresses (or not) in a stochastic manner [81, 82].

### Temporal correspondence between events in Gastruloids and embryos

The sequence of transcriptional and signalling events that we have described allows us to align the patterning of the *Gastruloids* with that of embryos and refine a previous estimate for the relative timing of events [13]. We surmise that the period between 48 and 72h AA corresponds to E5.5-E6.5 in embryos and serves to anchor our time-line (**Fig. 10**). This anchorage is corroborated by the patterns of expression of *Nodal*, *FGF4, 5, Wnt3* and *Wnt3a* that mirror the events in the embryo during this time period [83, 84] (**Fig. 3**). Furthermore, we observe re-expression of *Nanog* in the elongate at 72h AA (**Fig. 4E**), something that has also been described in the embryo in the nascent mesoderm at the start of Gastrulation [85]. It is in this period that AP polarity is firmly established, *T/Bra* expression becomes restricted to the proximal posterior region and gastrulation begins [82] and we see analogous events in the *Gastruloids* (see also van den Brink et al. [13]). The emergence of a node like structure at around 96h makes this time-point equivalent to E7.5 and the emergence of bilaterally asymmetric signalling associated with this structure shortly afterwards buttresses this comparison. Altogether these observations suggest a surprisingly close timing of events and of regulatory interactions between the *Gastruloids* and embryos that is summarised in **Fig. 10**.

**Figure 10:**
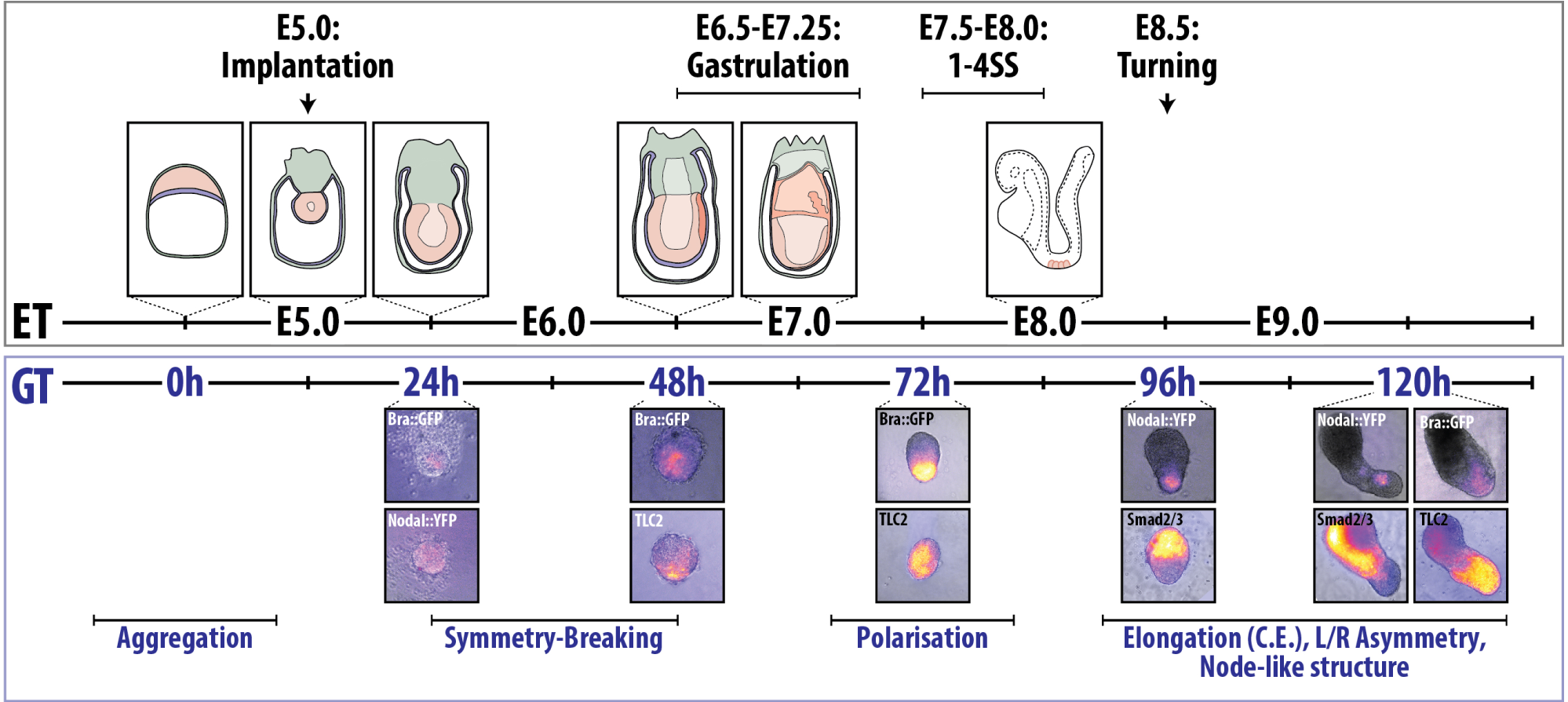
Alignment of the developmental stages of the mouse Embryo and *Gastruloids*. Top panel indicates *Embryo Time* (ET), displaying schematic representations of the mouse embryo with key developmental landmarks. Bottom panel indicates *Gastruloid Time* (GT) with images of *Gastruloids* at indicated time-points expressing the reporter constructs described in this study, aligned to the approximate developmental stage of the embryo (±0.5 days). Figure part adapted from [8] and the e-Mouse Atlas Project’s *Theiler Stages* website resource [62].

### Symmetry breaking *Gastruloids* and in mouse embryos

Our results show that between 24 and 48h AA, *Gastruloids* undergo a Nodal dependent symmetry-breaking event that, in collaboration with Wnt signalling, leads to the robust polarised expression of T/Bra::GFP to one pole which, by comparison with the embryo, we define as the posterior. We surmise that a similar symmetry-breaking event also occurs in the embryo around E5.5 but exploration of this possibility is constrained by the difficulties of working with embryos at this early stage. However, our system allows detailed analysis of the relationship between the different signalling and transcription factor networks which help enlighten these early stages. The suggested correspondence of events suggests that the period between 24 and 48h AA corresponds to between E4.5 and E5.5 and that this period is important for the establishment of the AP axis. While there is evidence for early engagement of Nodal in this process [42–44, 86], our results suggest an involvement of Wnt signalling in AP axis specification which has not been reported before. For example, inhibition of WNT signalling inhibition of Wnt signalling, e.g. with Porcupine mutants, shows no effect in the patterning of the embryo until the onset of gastrulation [87, 88]. This could be interpreted to suggest that the events that we observe in our system are not informative of the events *in vivo*. However, elements of Wnt signalling are expressed in the embryo from the blastocyst stage [89] and gain and loss of *β*-Catenin function, as well as of non canonical Wnt signalling, have effects earlier than those of Porcupine mutants [8]. Therefore, there might be a Wnt independent *β*-Catenin signalling event early in development that affects AP axis formation. We suggest that this is the case and that it is these activities that are revealed in our experiments. Our experimental system provides an opportunity to study this further.

Our results also highlight an important role for FGF signalling in the symmetry-breaking events. Inhibition of MEK signalling compromises the patterning of the *Gastruloids* in a manner similar to, but not as extreme as, that resulting from inhibition of Nodal signalling: loss of polarity, elongation and T/Bra::GFP expression. As in the case of Nodal signalling, this defect is not rescuable by Chi. On the other hand, exposure to FGF during 24-48h AA increases the domain of T/Bra::GFP expression and interferes, albeit temporarily, with elongation. In the embryo, it is often suggested that FGF signalling is required for the transition from the blastocyst to the epiblast [90–93] but analysis of *FGF4, 5, 8*, and FGF receptor mutants has not revealed any role in axis establishment though there are clear defects associated with mesoderm establishment and axial elongation [94–96]. Furthermore, exposure of early embryos to FGF leads to an expansion of the T/Bra::GFP expression domain [97] as we observe here (**Fig. 8**) and is consistent with a role in the onset of gastrulation.

As in the case of Wnt signalling, our results suggest a role of FGF signalling in the symmetry-breaking events which might also be operative but difficult to study in the embryo. Our study also shows that FGF/ERK signalling is closely associated with Wnt/*β*-Catenin signalling in the induction of *T/Bra* expression and therefore in the establishment of the Primitive Streak, and that this might be through modulation of the signalling event or through the target [98]. A related function of FGF in mesoderm induction has been described in other vertebrates [99, 100] but this is the first clear evidence that this might also be the case in mammals where it is often more closely associated with the transition from pluripotency to differentiation.

### Anteroposterior patterning in *Gastruloids* and embryos

A significant feature of the Wnt/*β*-Catenin driven patterning of the *Gastruloids* is the absence of proneural gene expression in the anterior domain [13]. When maintained in N2B27 many *Gastruloids* will develop *Sox1*-expressing cells characteristic of neural tissue [13, 14] and this default anterior neural fate is the basis for protocols to generate forebrain and eye cup structures [101–104]. However, after a pulse of Wnt/*β*-Catenin signalling, *Gastruloids* are able to generate spinal cord-like tissue because they are able to create Neuromesodermal progenitors, the pool that gives rise to this posterior structure and the paraxial mesoderm [14]. This independence of anterior and posterior development reflects the situation in the embryo where both regions are specified through separate mechanisms and, probably, at different times [105, 106]. This can be observed in, for example, loss of Dkk1 activity or in mutations resulting in a gain of *β*-Catenin [51, 107, 108] that produce embryos lacking anterior structures, reminiscent of *Gastruloids* after exposure to high levels of Wnt/*β*-Catenin signalling. It is noteworthy that gain of function constitutive mutations in the Wnt signalling receptor LRP6 or in *β*-Catenin, although extend the domain of Wnt signalling anteriorly, never cover the whole territory and leave a domain at the anterior most region of the embryo unresponsive [51]. This situation is also very similar to the one *Gastruloids* experience when they are exposed to high levels of Wnt signalling in which there is a transient ubiquitous response but in the anteriormost region the response is not stabilised.

Altogether, these observations suggest that when the AP axis is established the primitive streak specifies mesodermal and endodermal derivatives but that the anterior region of the embryo, that will give rise to the brain, is protected from this fate and delayed in its differentiation. The early suppression of BMP, Nodal and Wnt signalling does not lead to neural fate but rather to a pre-neural state, perhaps simply an epiblast state, protected from becoming primitive streak [109, 110]. This region becomes neural upon the release of antagonists of BMP and Nodal signalling from the prechordal plate and anterior definitive endoderm (ADE) at the end of gastrulation (reviewed in [111]). In a mirror image event, loss of BMP or Nodal signalling leads to embryos with, mostly, anterior neural structures [86, 112]. Failure to generate prechordal plate or the ADE leads to anterior truncations which can be compared to what we observe in our experiments [113, 114].

There is a clear and early subdivision of the aggregate into two domains that will respond differently to signals: Wnt/*β*-Catenin signalling during 48 and 72h AA stabilises the expression of T/Bra in the posterior region but it induces Nodal expression throughout the aggregate and, furthermore, there is a transient ubiquitous expression of T/Bra as well as of the Wnt/*β*-Catenin reporter expression that then relaxes to the pre-pulse situation. As transiently there are high levels of Nodal and Wnt/*β*-Catenin signalling in the anterior domain and there is no expression of T/Bra, this suggests that the symmetry-breaking event not only leads to a posterior like domain that will develop as in the embryo, but also that it generates an anterior domain that is refractory to this signalling event and upon which Wnt acts. The expression of high levels of the Smad2/3 reporter in this region confirm the increase in Nodal as well as highlight how this domain is refractory to *T/Bra* expression. On the other hand, this domain of Nodal signalling, provide an explanation for why this domain is also refractory to the neural fate as Nodal/Smad2/3 signalling will protect this domain from neural development. This suggests that *Gastruloids* have an anterior domain that is refractory to posterior development. It may be in this primed state and will be timely localised provision of BMP and Nodal inhibitors that releases this potential.

### Conclusions

Our results suggest that the symmetry-breaking event in the mouse embryo results from the self-organising activity of Nodal signalling around E5.0 which is amplified and stabilised at the proximal-posterior region of the embryo by interactions with agonists and antagonists of BMP, Wnt and Nodal itself, between E5.5 and E6.0. These interactions are patterned and lead to localisation of the initiation of gastrulation to a strategic point that allows the first cells that leave the Primitive Streak to invade the Extraembryonic Ectoderm. In the embryo it is likely that the threshold for this is raised by the extraembryonic tissue to ensure the role and outcome of the bias. Probably because of their characteristics, *Gastruloids* reveal and allow the amplification of this event. We have suggested a similar explanation of a related situation in the patterning of the neural tube in ESC-derived cysts [37]; perhaps there is a principle behind this which should be borne in mind in the development of tissues and organs in culture which may follow different paths than in embryos where their development will be constrained by the need to place them relative to other tissues and organs.

Our results show that the symmetry-breaking events in *Gastruloids*, an embryonic organoid system, recapitulate the main events in the embryo. Furthermore, their development closely mirrors that of embryos and allows us to position landmarks for an experimental time-line. Because of the ease of their manipulation, *Gastruloids* represent a good experimental system to explore the mechanisms underlying early developmental events. Here we have used them to reveal novel requirements and interactions for Wnt, Nodal and FGF signalling for symmetry-breaking that are extrapolatable to embryos.

There are other systems in which ESCs can be spatially patterned [115, 116] and each of them can make their own contribution to our understanding of the connection between cell fate assignments and the three dimensional organisation of tissues and organs. The observations that we report here on *Gastruloids*, particularly their ability to be cultured long term and the observation that in addition to an AP axis they can develop an LR asymmetry and generate a node-like structure, suggest that they might be useful beyond the early stages of development.

## Acknowledgements

We thank G. Keller for the T/Bra::GFP line, A-K Hadjantonakis for the TLC2 reporter and members of the AMA lab for useful discussions and criticisms. This work is funded by a European Research Council (ERC) Advanced Investigator Award to AMA (DAT, PCH) with the contribution of a Project Grant from the Wellcome Trust to AMA, an Engineering and Physical Sciences Research Council (EPSRC) Studentship to PB-J.

## 5 Supplemental Material

### 5.1 Generation of Spry4 reporter ES cell line

The Spry4 reporter construct was generated by combining ‘*knock-out first*’ targeting arms of the EUCOMM project [24] with a H2B-Venus reporter cassette and a neomycin resistance gene driven from a human *β*-actin promoter. This construct was integrated by homologous recombination into a cell line carrying a doxycycline-inducible Gata4-mCherry construct described in [25]. The construct used to obtain permanent expression of the H2B-Cerulean marker has been described [25]. To generate reporter lines, approximately 2×10^6^ cells were electroporated with 2 μg of the linearised targeting vector, plated onto feeder cells and put under selection one day after electroporation. Resistant colonies were picked after seven days, and PCR-genotyped for correct insertion events (reporter constructs) or visually screened for transgene expression. Karyotypes of the Spry4 reporter lines with the permanent nuclear marker were determined according to standard procedures [117], and clones with a median chromosome count of 40 were selected for experiments. Spry4::H2B-Venus reporter ESCs gave high-contribution chimaeras with germline transmission, confirming that this ES cell line retains full developmental potential.

### 5.2 Supplementary Figures

**Supplementary Figure 1:**
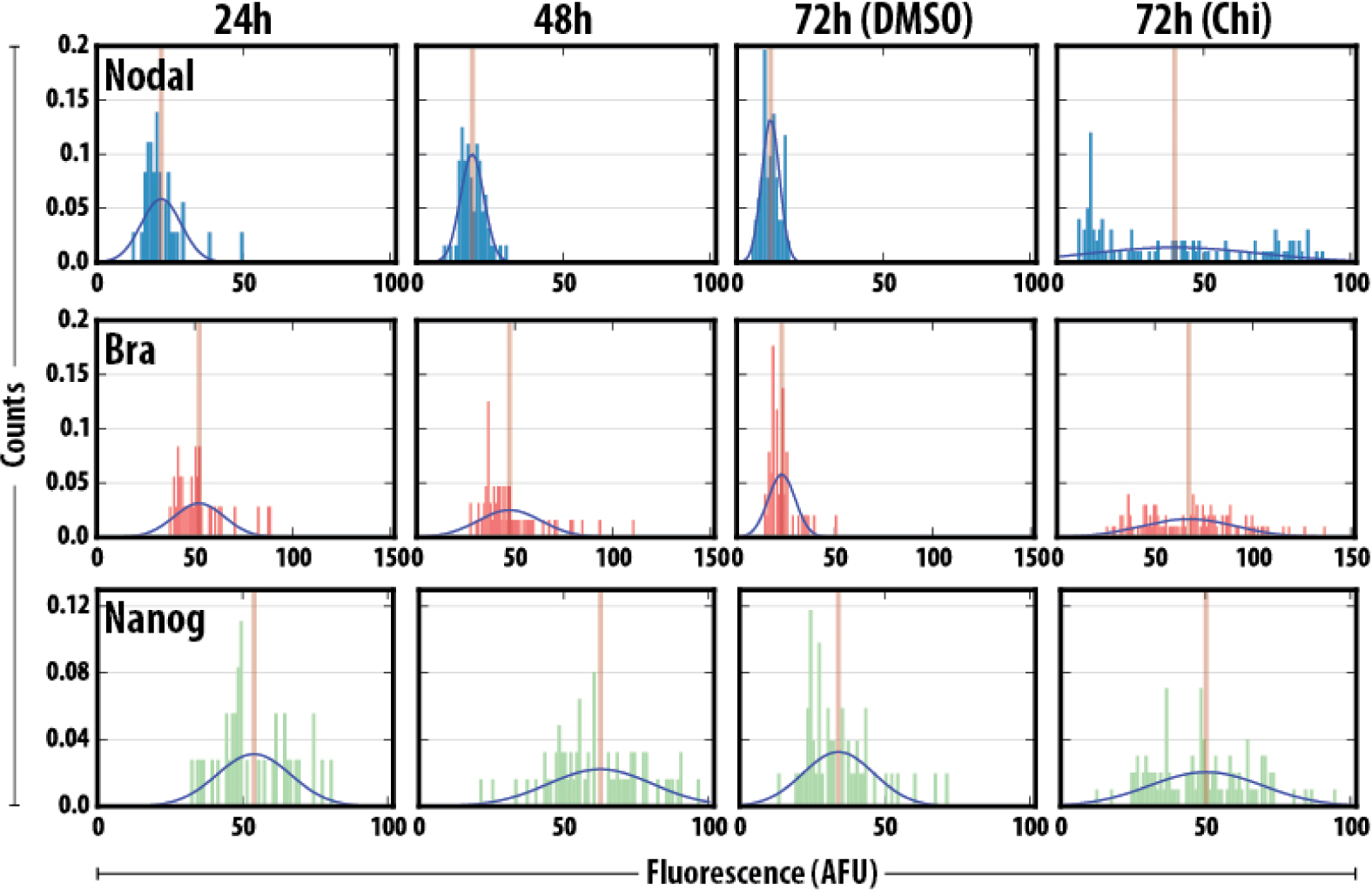
Quantification of T/Bra and Nanog expression in Nodal^condH BE::Y F P^ ESCs. A single *z*-plane from the confocal images of *Gastruloids* from **Fig. 4** was quantified and the distributions of YFP (Blue), T/Bra (Red) and Nanog (Green) plotted for the different treatments. The horizontal orange lines in each histogram correspond to the mean fluorescence levels.

**Supplementary Figure 2:**
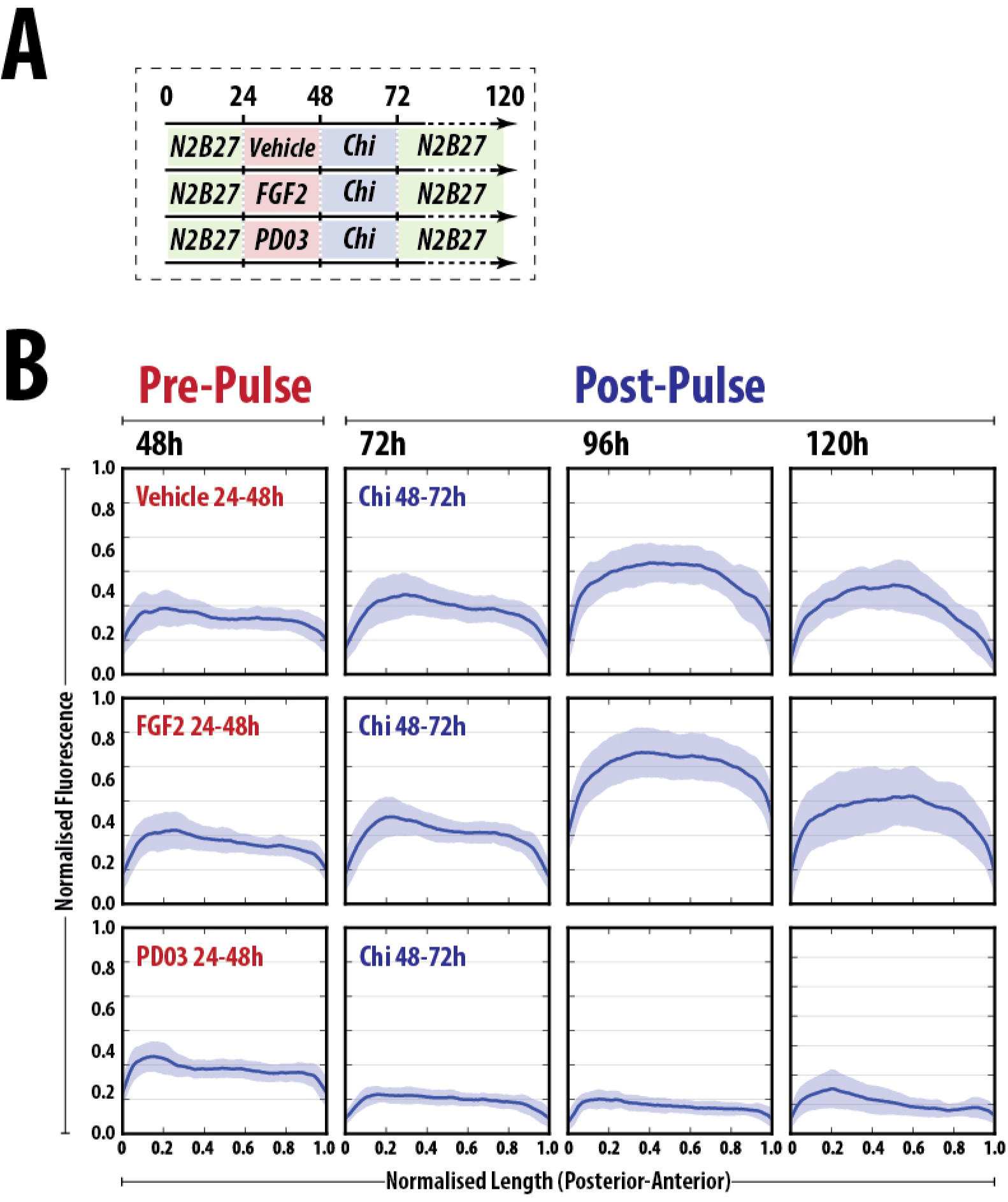
The activity of the Spry4::H2B-Venus reporter in *Gastruloids*. (A) Schematic representation of the experimental design. *Gastruloids* were aggregated in N2B27 (green shading) and stimulated pulsed with Chi between 48 and 72h AA following pre-treatment vehicle, FGF2 or PD03 between 24 and 48h AA. (B) The fluorescence levels of individual Spry4::H2B-Venus *Gastruloids* pre-treated as indicated (red text) were imaged by wide-field microscopy and quantitatively analysed for their fluorescence. The fluorescence plots were normalised to the maximum florescence of the control and the length of each *Gastruloid* was rescaled to have a maximum length of 1 unit.

**Supplementary Figure 3:**
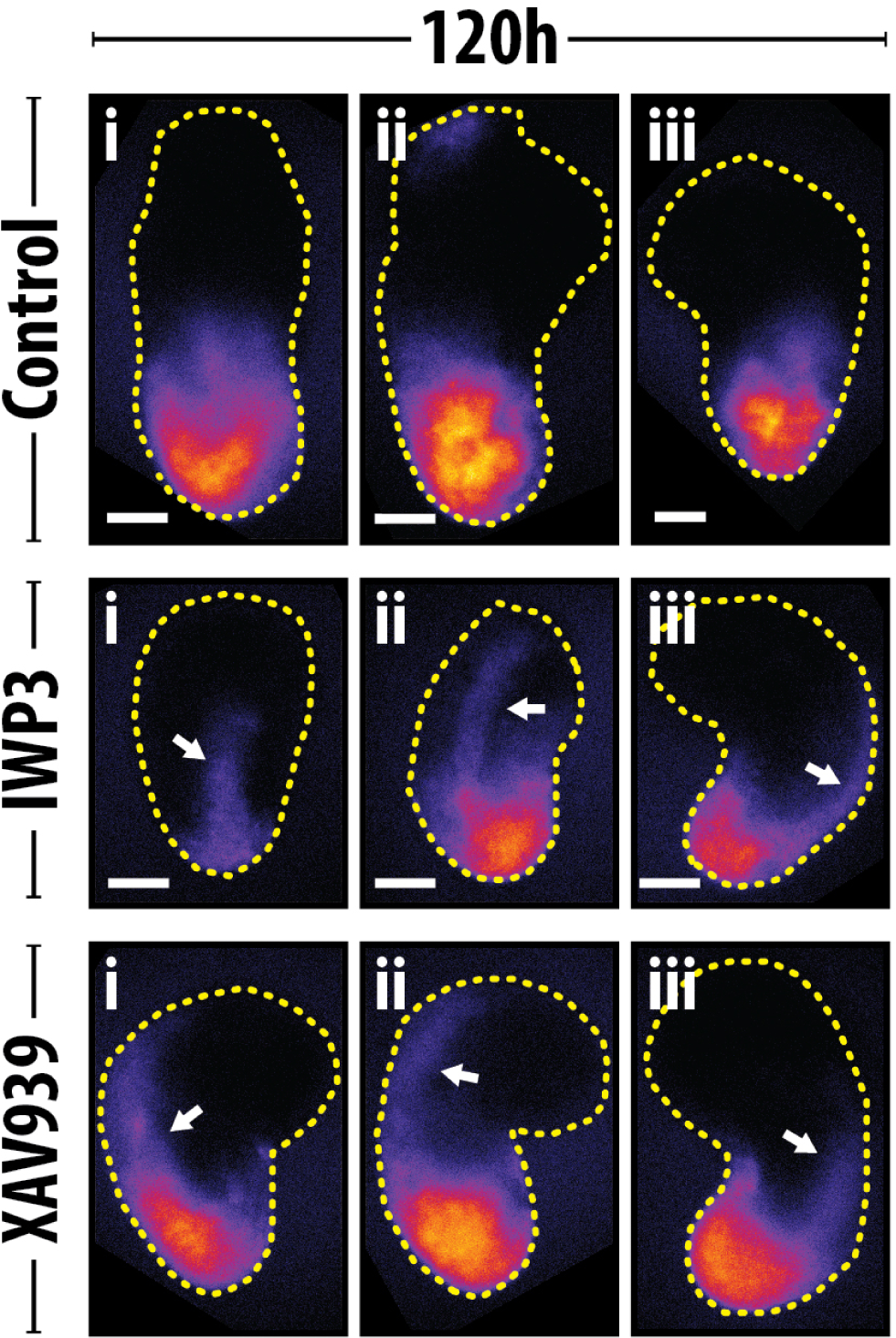
Inhibition of Wnt signalling prior to the Chi pulse results in midline expression in T/Bra::GFP *Gastruloids. Gastruloids* formed from T/Bra::GFP mouse ESCs were pre-treated with either DMSO (vehicle), IWP3 or XAV939 for 24h before wash-off and stimulation with Chi (see **Fig. 9A** for experimental set up); the 120h AA time-point is indicated. Notice the extended expression of T/Bra::GFP along the midline of the *Gastruloids* pre-treated with either IWP3 or XAV939 (indicated by white arrows). Three examples for each condition are shown. Scale bar represents 100 μm.

### 5.3 Supplemental Tables

**Table 1:** Antibodies used for *Gastruloid* immunostaining.

|  |  | Species | Dilution | Cat. Number | Supplier |
| --- | --- | --- | --- | --- | --- |
| Primary | Brachyury | Goat | 1:200 | sc-17743 | Santa Cruz Biotechnologies |
|  | Nanog | Mouse | 1:300 | 14-5761-80 | e-Biosciences |
|  | GFP | Rabbit | 1:500 | A11122 | Molecular Probes |
| Secondary | Goat-A633 | Donkey | 1:500 | A21082 | Molecular Probes |
|  | Mouse-A568 | Donkey | 1:500 | A10037 | Molecular Probes |
|  | Rabbit-A488 | Donkey | 1:500 | A21206 | Molecular Probes |
|  | Hoechst33342 | n/a | 1:1000 | H3570 | Invitrogen (ThermoFisher) |

**Table 2:** Cell lines used and numbers of cells required for *Gastruloid* culture. The average diameter of the *Gastruloids* at 48h post formation is indicated with the standard deviation and the number of *Gastruloids* measured from at least two replicate experiments.

| Cell Line | | Cells/40μl | 48h Diameter (μm) |
| --- | --- | --- | --- |
| AR8::mCherry | [18] | 550 | 182.7 ± 17.3 ($n = 83$) |
| Bra::GFP | [16] | 400 | 161.0 ± 26.2 ($n = 222$) |
| IBRE4::mCerulean | [18] | 400 | 152.6 ± 12.2 ($n = 39$) |
| Nodal<sup>condHBE::YFP</sup> | [17] | 500 | 138.7 ± 16.1 ($n = 124$) |
| Spry4::VNP | This work | 550 | 179.4 ± 20.3 ($n = 53$) |
| TLC2 | [19, 20] | 400 | 194.9 ± 20.7 ($n = 56$) |

**Table 3:** Concentrations of Small molecules and recombinant proteins used in this study. Conc: Concentration.

| Small Molecule/<br>Recombinant protein |  | Working Conc. | Stock Conc. | Supplier |
| --- | --- | --- | --- | --- |
| CHI99201 | [118] | 3μM | 10mM | Tocris Biosciences |
| PD0325901 | [119] | 1μM | 10mM | |
| SB431542 | [120] | 10μM | 100mM | |
| DMH1 | [121, 122] | 0.5nM | 5mM |  |
| IWP3 | [35] | 1μM | 5mM | |
| XAV939 | [36] | 1μM | 10mM | |
| Wnt3a | | 100ng/ml | 40μg/ml | R&D systems |
| DKK | | 200ng/ml | 100μg/ml | |
| Nodal | | 1μg/ml | 50μg/ml | |
| Noggin | | 500ng/ml | 100μg/ml | |
| Chordin | | 500ng/ml | 200μg/ml | |
| FGF2 | | 10ng/ml | 100μg/ml | |

**Table 4:**
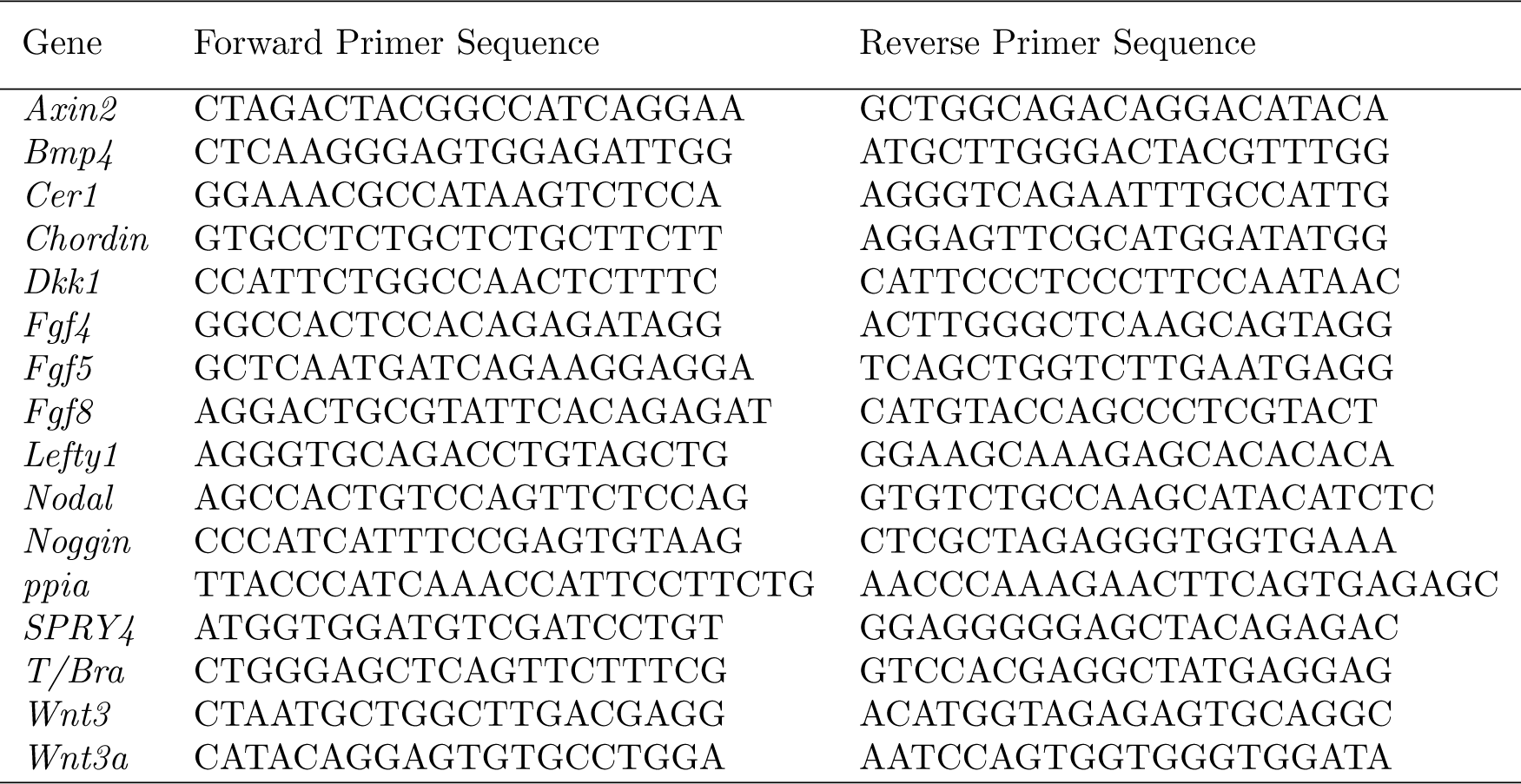
Primer Sequences used for qRT-PCR.

**Table 5:**
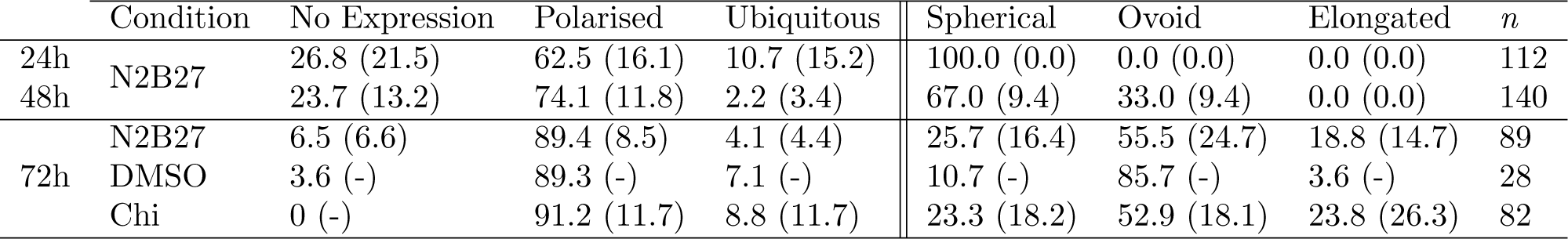
Expression phenotype of T/Bra::GFP mouse ESCs. The proportion of T/Bra::GFP *Gastruloids* not expressing the reporter (No Expression) or displaying either Polarised or Ubiquitous expression at 24, 48 and 72h AA when maintained in N2B27 (24, 48, 72h) or following a pulse of DMSO or Chi (72h). The standard deviation is shown in brackets and the number of *Gastruloids* analysed are shown (*n*).

**Table 6:** Pearson Correlation Coefficients between a) Nodal and Brachyury, b) Nodal and Nanog and c) Brachyury and Nanog in the Nodal^condH BE::Y F P^ mouse ESC line at 24, 48 and 72h with or without a pulse of Chi on day 3. Values to 3d.p., correlations from one representative *Gastruloid*.

|  | Pearson Correlation Coefficient |  |  |  |
| --- | --- | --- | --- | --- |
|  | 24h | 48h | N2B27 72h | Chi 72h |
| a) Nodal:Brachyury | -0.029 | 0.592 | 0.807 | 0.643 |
| b) Nodal:Nanog | 0.112 | 0.452 | 0.814 | 0.280 |
| c) T/Bra:Nanog | 0.391 | 0.443 | 0.731 | 0.327 |

